# Pancreatic cancer ductal cell of origin drives CD73-dependent generation of immunosuppressive adenosine

**DOI:** 10.1101/2021.11.29.470415

**Authors:** Kanchan Singh, Erika Y. Faraoni, Yulin Dai, Vidhi Chandra, Emily Vucic, Tingting Mills, Melissa Pruski, Trent Clark, George Van Buren, Nirav C. Thosani, John S. Bynon, Curtis J. Wray, Dafna Bar-Sagi, Kyle L. Poulsen, Lana A. Vornik, Michelle I. Savage, Shizuko Sei, Altaf Mohammed, Zhongming Zhao, Holger K. Eltzschig, Powel H. Brown, Florencia McAllister, Jennifer Bailey-Lundberg

**Affiliations:** Department of Anesthesiology, McGovern Medical School, The University of Texas Health Science Center at Houston, Houston, TX 77030; Center for Precision Health, School of Biomedical Informatics, The University of Texas Health Science Center at Houston, Houston, TX 77030; Department of Clinical Cancer Prevention, The University of Texas MDAnderson Cancer Center, Houston, TX 77030; Departments of Biochemistry and Molecular Pharmacology and Medicine, NYU Langone School of Medicine, New York, NY 10016; Department of Biochemistry, McGovern Medical School, The University of Texas Health Science Center at Houston, Houston, TX 77030; Division of Gastroenterology, Hepatology and Nutrition, Department of Internal Medicine, McGovern Medical School, The University of Texas Health Science Center at Houston, Houston, TX 77030; Division of Surgical Oncology, Baylor College of Medicine, Houston, TX, 77030; Center for Interventional Gastroenterology at UTHealth (iGUT), McGovern Medical School, The University of Texas Health Science Center at Houston, Houston, TX 77030; Department of Surgery, McGovern Medical School, The University of Texas Health Science Center at Houston, Houston, TX 77030; Center for Perioperative Medicine, Department of Anesthesiology, McGovern Medical School, The University of Texas Health Science Center at Houston, Houston, TX 77030; Division of Cancer Prevention, National Cancer Institute, Rockville, MD 20850; Department of Gastrointestinal Medical Oncology, The University of Texas MDAnderson Cancer Center, Houston, TX 77030; Department of Immunology, The University of Texas MDAnderson Cancer Center, Houston, TX 77030

**Keywords:** Cell of origin, immunosuppression, adenosine, pancreatic cancer subtypes

## Abstract

The microenvironment that surrounds pancreatic ductal adenocarcinoma (PDAC) is profoundly desmoplastic and immunosuppressive. Understanding initial triggers of immunosuppression during the process of pancreatic tumorigenesis would aid in establishing novel targets for effective prevention and therapy. Here, we interrogate the differential molecular mechanisms dependent on cell of origin and pathology subtype that determine immunosuppression during PDAC initiation and in established tumors. Transcriptomic analysis of cell of origin dependent-epithelial gene signatures revealed that *Nt5e*/CD73, a cell surface enzyme that is the pacemaker for extracellular adenosine generation, is one of the top 10% of genes over-expressed in murine tumors arising from ductal pancreatic epithelium as opposed to those rising from acinar cells. These findings were confirmed by Imaging Mass Cytometry and High-Performance Liquid Chromatography. Our data indicate that ductal activation of oncogenic mutant *Kras* results in loss of PTEN and elevated AKT signaling which ultimately releases CD73 suppression. Delivery of CD73 small molecule inhibitors through various delivery routes reduced tumor development and growth in genetically engineered and syngeneic mouse models. Analysis in human PDAC subtypes indicates that high *Nt5e* in murine ductal PDAC models overlaps with high *NT5E* in human PDAC Squamous and Basal Subtypes, considered to have the highest immunosuppression and worst prognosis. These findings highlight a molecular trigger of the immunosuppressive PDAC microenvironment which is dependent on ductal cell of origin, linking biology with pathological subtype classification, critical components to personalized approaches for PDAC prevention and immunotherapeutic intervention.

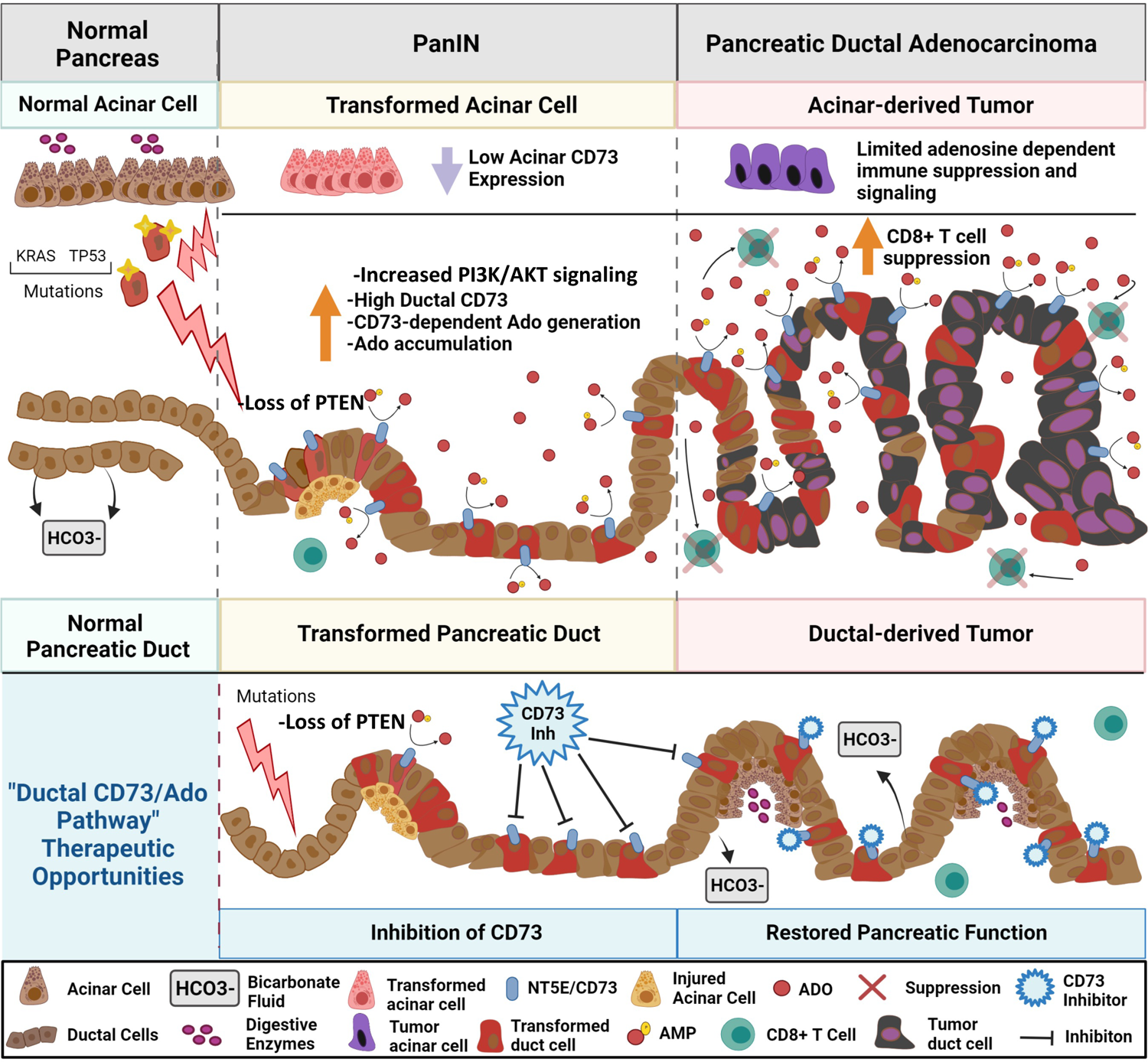

## INTRODUCTION

Pancreatic ductal adenocarcinoma (PDAC) remains very deadly and is predicted to become the second leading cause of cancer-related deaths by the year 2030 in the United States (1, 2). Clinical trials for PDAC aim to target various components of the complex fibrotic immunosuppressive tumor microenvironment (3–6). CD73 is a key enzyme involved in dampening inflammatory responses in inflamed and hypoxic tumor microenvironments (7). CD73 is an ectoenzyme that can be soluble, but is predominantly localized on the cell surface and generates extracellular adenosine. Extracellular adenosine is an important anti-inflammatory nucleoside involved in the resolution stage of inflammation and tissue repair by signaling through adenosine receptors (8). In the absence of CD73 in inflammatory conditions, increased extracellular adenosine triphosphate (ATP) mediates inflammation and purinergic signaling by binding to purinergic receptors. In the presence of ectonucleotidase triphosphate diphosphohydrolase-1 (CD39), a cell surface enzyme with catalytic activity, ATP is rapidly converted to AMP, which is the substrate for catalytic conversion to immunosuppressive adenosine by CD73.

In recent years, adenosine has been shown to have profound immunosuppressive and pro-angiogenic effects in the tumor microenvironment and a number of pre-clinical models have shown targeting CD73 has potent anti-tumor effects (9–12). However, its regulators, role in generating adenosine and modulating tumor progression in pancreatic cancer remains undefined. Previous studies have shown the role and expression of CD73 can deviate between subtypes of tumors as well as cell types of the tumor microenvironment(13, 14).

In this manuscript, we aimed to identify stromal changes and therapeutic vulnerabilities based on cell of origin, comparing acinar-derived vs ductal-derived murine PDAC whole transcriptomic signatures. We found that ductal derived PDAC had pathway enrichment implicating a strong immunosuppressive microenvironment. The top Gene Ontology (GO) category identified from RNA-seq signatures in acinar and ductal derived PDAC was leukocyte cell-cell adhesion. We observed *Nt5e*/CD73 was one of the top 5 genes most significantly elevated in ductal compared to acinar PDAC in this GO category.

Subtypes of human PDAC have been defined using comprehensive whole transcriptomic profiling of large cohorts of patients (1, 15, 16). Human PDAC subtypes have differences in somatic mutations which may also dictate response to chemotherapy or immunotherapy and predict overall survival (15, 17). The transcriptomic signatures from tumors have been widely used for cancer type classification and prognosis(18, 19). To determine if murine cell of origin models compared to human PDAC subtypes, we compared cell of origin whole transcriptomic signatures from mouse models to those of published human subtypes. In addition to sequencing data, we analyzed early alterations in the tumor microenvironment in ductal-derived Pancreatic intraepithelial neoplasia (PanIN) and PDAC. While somewhat controversial, emerging data has shown overall tumor mutational burden (TMB) may constitute a biomarker for response to immunotherapy in some solid malignancies(20–24). However, PDAC, patients with high tumor mutational burden have not shown higher response rates to anti-PD-1/PD-L1 immunotherapy, highlighting that PDAC biology is very unique and that concepts cannot be easily translated from other cancer to PDAC (25, 26).

## RESULTS

### Comparison of ductal and acinar cell derived tumor signatures to human molecular subtypes of PDAC

Using the *Hnf1b:CreERT2* (*C^Duct^*) ductal specific inducible allele or a *Ptf1a:CreERTM* (*C^Acinar^*) acinar cell specific allele, we generated genetically engineered mouse models (GEM) to study cancer development from ductal or acinar cells. Inducible CreER mice were crossed to *LSL-Kras^G12D^* and *LSL-Trp53^R172H^* mice to generate KPC^Duct^ and KPC^Acinar^ GEM models (**Figure 1A-B**). As we and others have shown, both KPC^Duct^ and KPC^Acinar^ GEM models develop moderately to well differentiated PDAC (**Figure 1A-B**)(27–42). To determine if KPC^Duct^ or KPC^Acinar^ tumor signatures are enriched in published human PDAC whole transcriptomic subtypes(16, 17, 43), Gene Set Variation Analysis (GSVA) was used to analyze and compare our murine gene signatures with human molecular subtypes previously defined by the Australian International Cancer Genome Initiative (ICGC) and The Cancer Genome Atlas (TCGA) Research Network. We defined KPC^Acinar^ and KPC^Duct^ tumor signatures based on their highest-confidence differentially expressed genes (Methods). The ortholog genes between human and mouse ductal and acinar signatures revealed a significant difference in ductal-derived and acinar-derived whole transcriptomic profiles, with a ductal-derived tumor signature significantly enriched *(**P=0.0075)* in correlated Squamous human PDAC tumors and the murine acinar-derived tumor signature was significantly enriched *(*P=0.014)* in Immunogenic PDAC tumors (**Figure 1C-E**). In addition to Immunogenic and Squamous subtypes, we compared our murine PDAC signature genes to the Cancer Genome Atlas (TCGA) Basal and Classical RNA-Seq profiles. Heatmap and violin plot analysis revealed the ductal-derived tumor signature significantly correlated *(***P=0.0051)* with Basal human PDAC tumors and acinar-derived tumor signature correlated with Classical subtype (**Figure 1F-H**). Our analysis revealed murine tumors arising in Hnf1b+ pancreatic ducts have RNA-seq signatures that significantly overlap with the Squamous and Basal human PDAC subtypes while acinar derived tumors have RNA-seq signatures that align with Immunogenic and Classical subtypes.

**Figure 1.**
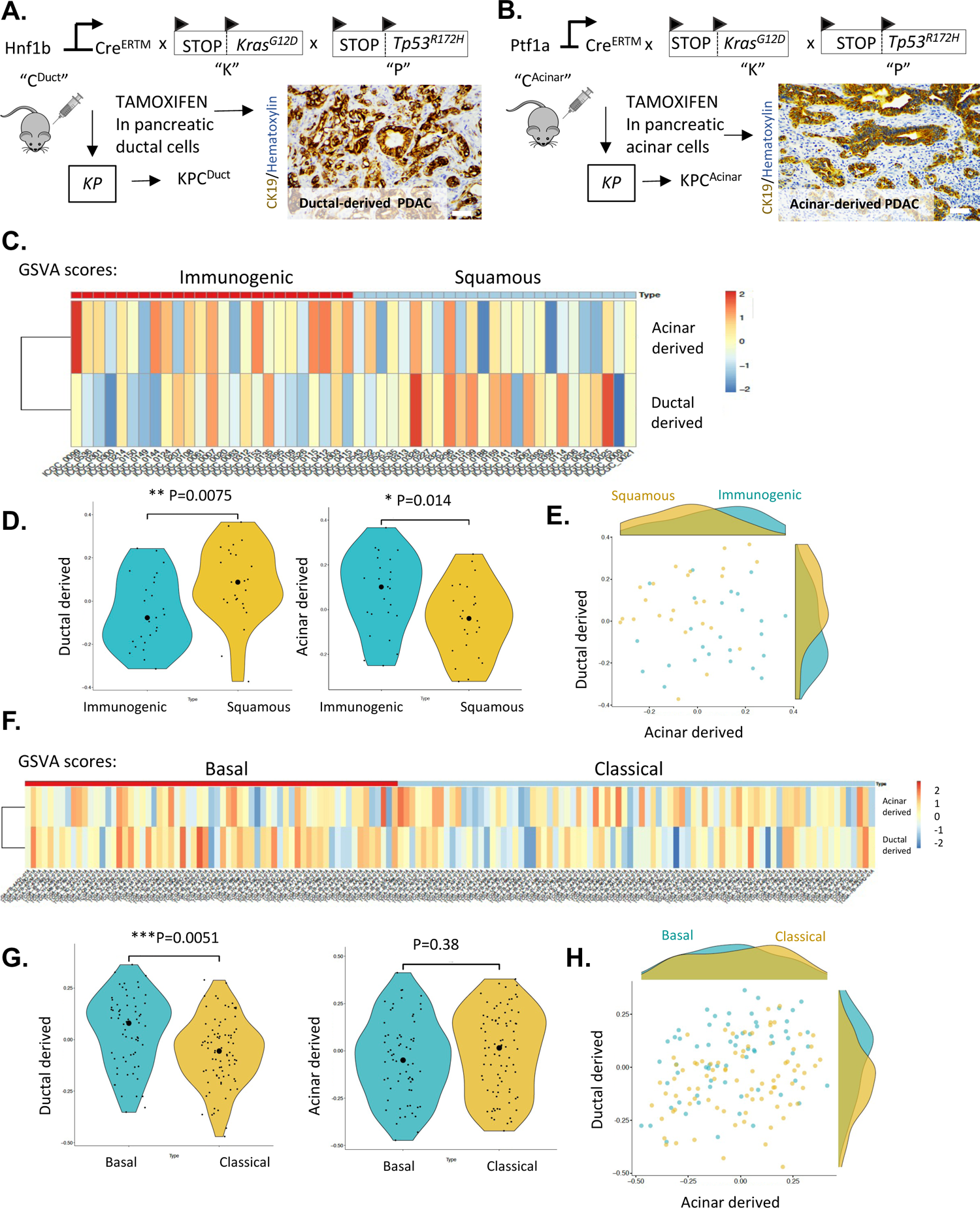
Comparison of ductal and acinar cell derived tumor signatures to human molecular subtypes of PDAC. **A-B)** Schematic of transgenic mouse breeding scheme to generate mutant *Kras* and *Tp53* tumors from acinar and ductal cells and IHC analysis of CK19 to show ductal adenocarcinoma arising in ductal and acinar mouse models of PDAC. **C)** Heatmap shows the GSVA scores for human homolog genes derived from mouse ductal and acinar signatures in different samples from ICGC immunogenic and squamous subtypes. The color represents the relative GSVA score. **D)** Violin plot of the GSVA scores for human homolog genes derived from mouse ductal and acinar signature genes in ICGC immunogenic and squamous subtypes. **E)** Scatter plot of the GSVA scores for human homolog genes derived from mouse ductal and acinar signature genes in ICGC immunogenic and squamous subtypes. **F)** Heatmap shows the GSVA scores for human homolog genes derived from mouse ductal and acinar signatures in different samples from TCGA Basal and Classical subtypes. **G)** Violin plot of the GSVA scores for human homolog genes derived from mouse ductal and acinar signature genes in TCGA Basal and Classical subtypes. **H)** Scatter plot of the GSVA scores for human homolog genes derived from mouse ductal and acinar signature genes in TCGA Basal and Classical subtypes. The big round dots represent the medium of the GSVA scores, while the small round dots represent the score for each sample. We used a non-parametric Wilcoxon rank sum test for both groups. * indicates p < 0.05; ** indicates p < 0.01. These data reveal murine tumors arising in pancreatic ducts have RNA-seq signatures that significantly overlap with the Squamous and Basal human PDAC subtypes while acinar derived tumors have RNA-seq signatures that align with Immunogenic and Classical Subtypes.

### Ductal-derived tumors have a distinct immunosuppressive gene signature

We used Ingenuity Pathway Analysis (IPA) to examine canonical pathway analysis of murine KPC^Duct^ and KPC^Acinar^ RNA-seq signatures. Principal component analysis (PCA) and Volcano plots show distinct gene clustering of acinar-derived versus ductal-derived PDAC (**Figure 2A,B**). IPA analysis of Top Altered Pathways increased (orange) or decreased (blue) in murine cell of origin tumors (**Figure 2C**) revealed an immunosuppressive signature in ductal-derived tumors with Immune Response to leukocytes, T cell response, Migration of myeloid cells and T cell development all significantly downregulated in ductal-derived murine PDAC compared to acinar-derived PDAC. Further analysis of Top Diseases and Functions revealed significantly down-regulated IPA pathways in ductal-derived tumors were Crosstalk between Dendritic Cells and Natural Killer Cells, Th1 Pathway, Natural Killer Cell Signaling and Inflammasome Pathway, which reinforced an immunosuppressive signature (**Supplemental Figure 1**), In addition to IPA, we evaluated our differentially expressed genes using GO to define top enriched function categories. Leukocyte cell-cell adhesion was the highest GO category (Benjamini and Hochberg procedure corrected p_BH_ = 0.0004) identified (**Figure 2D**) and Nt5e was significantly elevated in this category (**Figure 2E**; **Supplemental Table 1**). Given the higher *Nt5e*/CD73 expression in KPC^Duct^ compared to KPC^Acinar^, we examined if *NT5E*/CD73 was also differentially expressed in previously published human Squamous vs. Immunogenic and Basal vs. Classical subtype comparisons (16, 17). Notably, *NT5E*/CD73 was also found elevated in human Squamous and Basal subtypes relative to Immunogenic and Classical, placing *NT5E*/CD73 as one of the 21 highly expressed overlapping genes between murine KPC^Duct^ and human Squamous and Basal Subtypes, as shown by Venn diagram analysis (**Figure 2F-G**; Supplemental Table 2).

**Figure 2.**
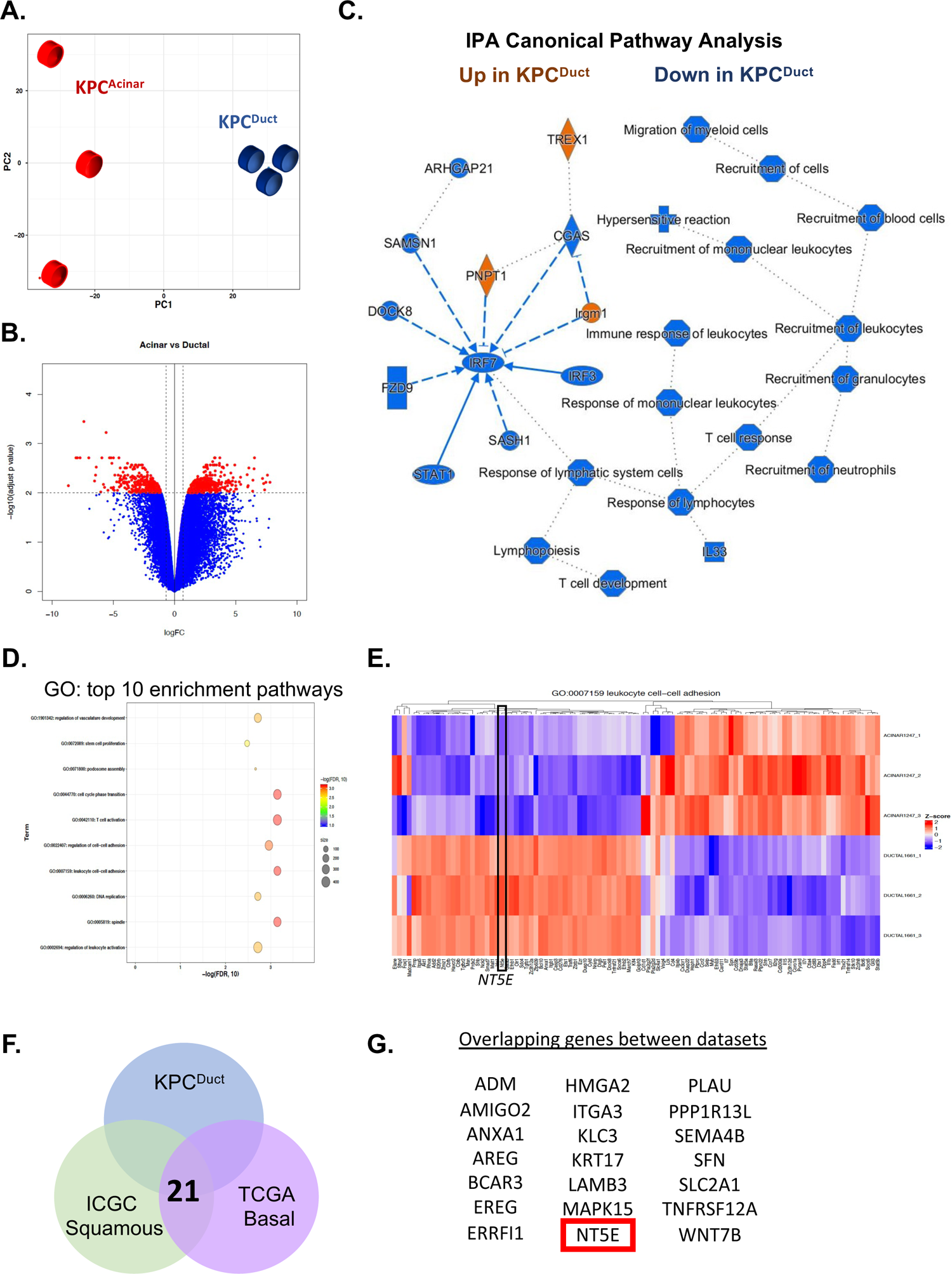
*NT5E/CD73* is highly expressed in murine ductal derived PDAC and high CD73 expression correlates with Basal and Squamous human PDAC subtypes. **A)** PCA analysis of RNA-seq samples. Tumors arising in ductal and acinar cells have distinct profiles. **B)** Volcano plot showing significantly expressed genes in acinar versus ductal derived PDAC**. C**) IPA analysis of Top Altered Pathways increased (orange) or decreased (blue) in murine cell of origin tumors. **D)** Gene ontology top 10 enrichment pathways altered in acinar-derived versus ductal-derived PDAC. **E)** Leukocyte cell-cell adhesion was tone of the top differentially expressed GO categories and *nt5e* was a significantly elevated gene in ductal-derved PDAC in this category. **F)** Venn diagram showing the number of top overlapping genes in murine KPC^Duct^ and human Squamous and Basal Subtypes. **G)** NT5E/CD73 is one of the top overlapping genes expressed in all the data sets.

### CD73 is over-expressed in murine and human ductal-derived PanINs and PDAC

To directly compare stromal and epithelial mechanisms of ductal vs. acinar-derived *Kras* dependent development of PanIN, we crossed the *C^Acinar^* allele to the *Kras^G12V^* allele to generate KC^Acinar^ mice (**Figure 3A**). Similar to what we recently observed and published in pancreatic ducts expressing *Kras^G12V^* (KC^Duct^), (**Figure 3A**), tamoxifen given at a moderate dose (5mg) resulted in acinar-derived PanIN, fibrosis and PDAC (**Figure 3B**; **Supplemental Figure 2**). We then observed abundant CD73 protein expression in ductal, but not acinar-derived PanIN epithelium (**Figure 3B**). Furthermore, we also confirmed CD73 over-expression on the apical luminal side in KPC^Duct^ but not KPC^Acinar^ tumors (**Figure 3C**). As we observed significantly elevated NT5E in human PDAC subtypes associated with poor prognosis, we wanted to determine if *NT5E* levels correlated with poor prognosis in human PDAC. Analysis of TCGA data revealed significantly elevated *NT5E* is a predictor of poor overall survival in patients with PDAC (**Figure 3D**).

**Figure 3.**
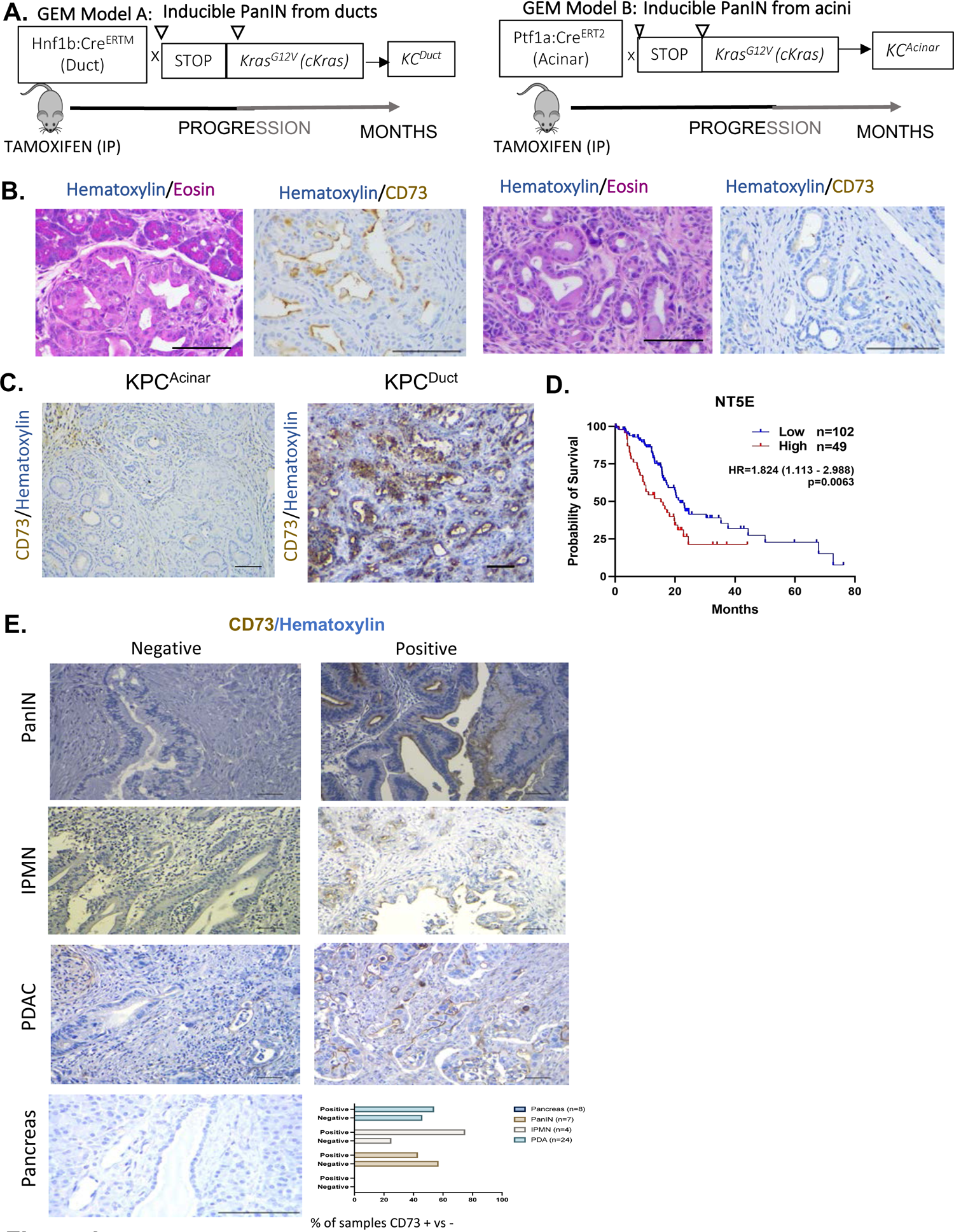
CD73 expressed in murine and human PDAC generates immunosuppressive adenosine. **A)** Schematic of mouse models to generate PanIN and PDAC from ductal or acinar cells using an inducible *Kras^G12V^* allele**. B)** Representative histology of PanIN arising in ductal epithelium and CD73 IHC showing high expression of CD73 in ductal derived PanIN and lack of staining for CD73 in acinar derived PanIN. **C)** IHC analysis of CD73 expression in acinar and ductal derived murine PDAC. **D)** TCGA analysis reveals high expression of *NT5E*/CD73 in human PDAC is significantly correlates with worse prognosis. **E)** Immunohistochemical labeling for CD73 in a human PDAC tissue array. CD73 is expressed in 54% of PDAC histologic subtypes and 75% of malignant IPMN (scale bars are 50um).

As we observed predominantly epithelial labeling of CD73 in our murine ductal-derived PanIN and PDAC samples, we wanted to determine the cellularity of CD73 in human PDAC, PanIN and intraductal pancreatic mucinous neoplasia (IPMN), another ductal precursor lesion. While we did not observe staining for CD73 in normal pancreas, we observed epithelial CD73 expression in neoplastic epithelium (42% of PanIN (n=12) and 75% of malignant IPMN (n=4)) and 54% of PDAC we analyzed (n=44) (**Figure 3E**).

### Ductal-specific CD73 over-expression results in development of adenosine

To determine if elevated CD73 resulted in increased parenchymal adenosine, we evaluated intrapancreatic adenosine levels from KC^Duct^ and KC^Acinar^ mice by High-Performance Liquid Chromatography (HPLC). We observed significantly elevated adenosine levels in KC^Duct^ pancreata compared to wild type or KC^Acinar^ pancreata (**Figure 4A**). Notably, we also found that levels of adenosine monophosphate (AMP), the substrate for CD73, were significantly increased in wild type and KC^Acinar^ pancreata compared to KC^Duct^ (**Figure 4B**).

**Figure 4.**
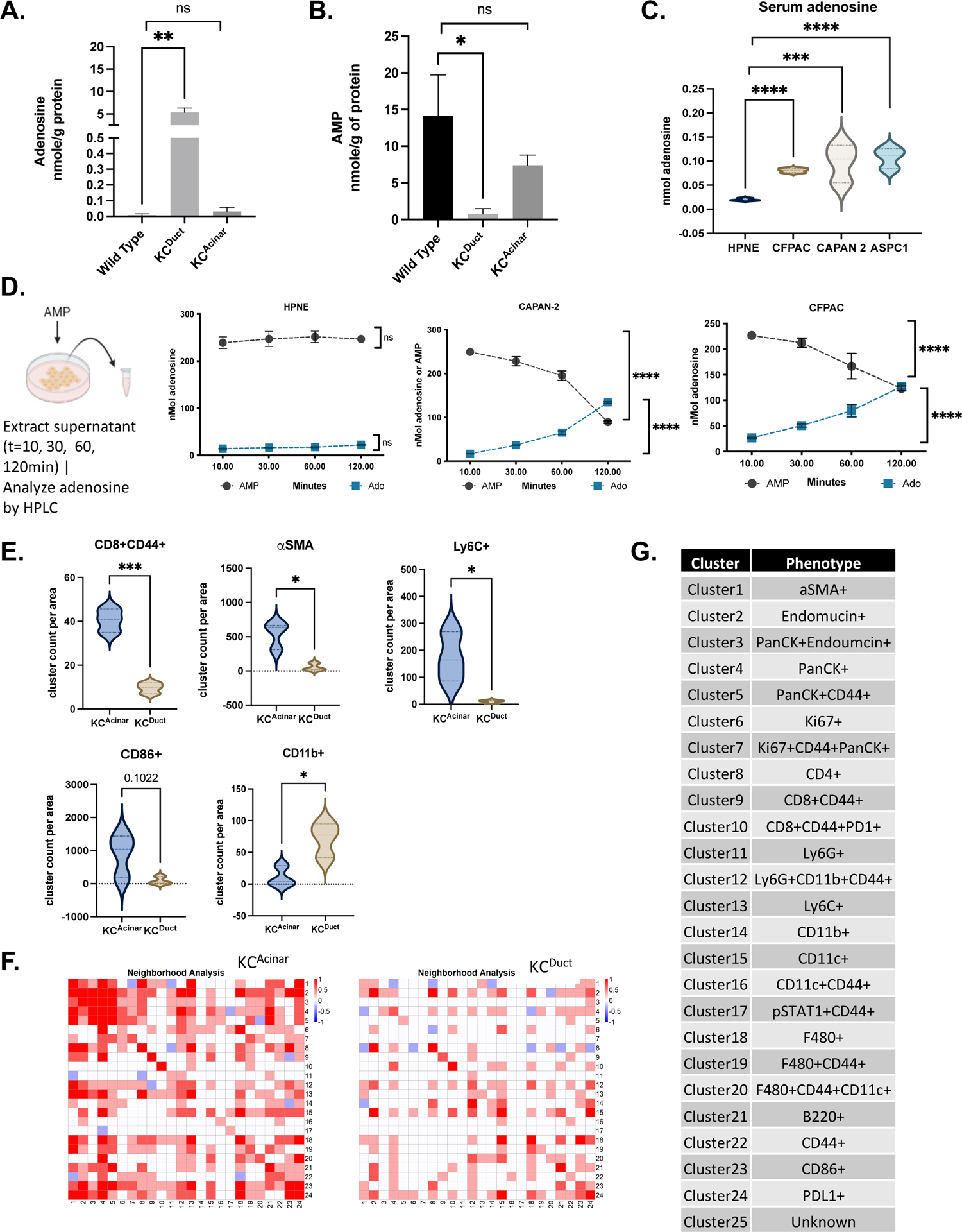
Imaging Mass Cytometry (IMC) profiling reveals an immunosuppressive environment in ductal derived PDAC. **A**) HPLC analysis of adenosine and **(B)** AMP levels in wild type pancreas, KC^Duct^ or KC^Acinar^ N=3 samples per group. Intrapancreatic adenosine levels are significantly elevated in KC^Duct^ and pancreata consistent with CD73 expression in KC^Duct^ neoplasia *(**P<0.01)* and intrapancreatic AMP levels are highest in wild type and KC^Acinar^ tissue indicating lack of CD73 activity in these tissues (*\*P<0.05*). **C)** Serum adenosine levels are significantly increased in PDAC cell lines relative to control pancreatic cells (HPNE) *(P<0.001; P<0.0001*), a student’s t-test was used for statistical comparison. **D)** To functionally evaluate and measure CD73 activity in cell lines *in vitro*, AMP (250ng) was administered to fresh HBSS (supernatant) and samples were taken at 10, 30, 60 and 120 minutes to measure AMP and adenosine levels. We observed a significant increase in adenosine generation in human PDAC cells (CAPAN-2, CFPAC) compared to control HPNE cells *(****P<0.0001*; a student’s t-test was used for statistical analysis). **E)** Quantitative clustering analysis reveals CD8+CD44+ T cells, aSMA+ cells, Ly6C+, and CD86+ cells are increased in acinar versus ductal derived PanIN and PDAC *(***P<0.001*). Graphical representation of immune and stromal clusters evaluated by IMC. *P* values were calculated using a student’s t-test in Prism Graphpad Software *(*P<0.05); (***P<0.001).* **F)** Neighborhood analysis IMC data show significant association of activated CD8 T cells near PanCK cells in KC^Acinar^ but not KC^Duct^ pancreata. **G)** IMC cluster identification.

To assess CD73 activity, we analyzed adenosine levels in the supernatant of human and murine PDAC cell lines compared to normal human HPNE pancreatic cells(44). We observed significantly elevated adenosine in CFPAC, CAPAN-2 and ASPC1 supernatant compared to normal human (HPNE) cells (**Figure 4C**). Supernatant was analyzed by HPLC and we observed a significant decrease in serum AMP levels in all human PDAC cell lines and a concomitant significant increase in serum levels of adenosine (**Figure 4D**). In contrast, we did not observe a significant increase in adenosine levels in HPNE cells indicating high CD73 activity is unique to PDAC cells.

### Ductal-derived PanIN and PDAC generate an immunosuppressive tumor microenvironment

As we observed differences in CD73 epithelial staining, we sought to comprehensively immunoprofile acinar- and ductal-derived microenvironments through comprehensive imaging mass cytometry (IMC) (**Figure 4E**; **Supplemental Figure 3**). Dimension reduction from IMC analysis revealed striking immunoprofiling differences in numbers of CD8a+ cells per area from KC^Acinar^ and KC^Duct^ pancreata. IMC profiling revealed significantly increased activated CD8+CD44+ T cells, Ly6C+ and elevated CD86+ cells (**Figure 4E**) in KC^Acinar^ tissue compared to KC^Duct^ tissue. We also observed significantly increased αSMA in acinar-derived PanIN and PDAC sections, similar to what has recently been described in murine acinar-derived compared to ductal-derived PDAC(28). We then generated neighborhood analysis to determine spatial differences in immune cell subtypes in our GEM models. We observed activated CD8+ T cells are adjacent to PanCK cells in KC^Acinar^ but are absent in KC^Duct^ pancreata (**Figure 4F**). Neighborhood analysis also revealed Ly6C+ and CD11c+ cells adjacent to PanCK epithelium in KC^Acinar^ pancreata. In contrast, we observed increased number of PD-1+ CD8+ T cells adjacent to PanCK+ epithelium in the KC^Duct^. While activated CD8+ T cells were not observed near KC^Duct^ PanCK+ cells, CD86+, F4/80+ and CD11c+ cells were observed in moderate proximity to PanCK+ epithelium (**Figure 4F**).

### Mutant *Kras* elevates levels of *NT5E* in pancreatic ducts

We recently published a GEM model of ductal-derived PanIN and PDAC. In this model system, we published recombination of *Kras^G12V^* in 50% of pancreatic ducts resulted in total K-Ras levels slightly less, but comparable to, those observed in KPC^Duct^ mice(35) (**Supplemental Figure 4A,B**). Notably, while K-Ras protein was comparable, Ras-GTP levels were 2-10-fold higher in KPC^Duct^ mice compared to KC^Duct^ GEM (**Supplemental Figure 4A,B**). We hypothesized based on CD73 positive staining for in KPC^Duct^ and KC^Duct^ pancreata (**Supplemental Figure 4C**) that mutations in *Kras* alone are important for elevating levels of ductal CD73. Thus, we examined early molecular alterations in *Kras^G12V^* ducts to study if oncogenic Kras alone could elevate levels of CD73. To test our hypothesis, we administered 0 (Tam^0^) or 10mg (Tam^10^) of tamoxifen to generate ductal expression of mutant *Kras^G12V^* levels *in vivo* (**Figure 5A**). We then waited four days before euthanizing KC^Duct^ GEM mice. After euthanasia, ductal cells were cultured *ex vivo* as we have previously described and after seven days RNA was extracted for RNA-Seq (**Figure 5B**). Volcano plot analysis from differentially expressed genes revealed divergent gene signatures in Tam^0^ compared to Tam^10^ ducts (**Figure 5C**). In addition, GO analysis on differentially upregulated genes between Tam^10^ and Tam^0^ ducts revealed major GO categories elevated in Tam^10^ compared to Tam^0^ ducts including Regulation of Inflammatory Response (p_BH_ = 6.43 × 10^-6^), which contains *Nt5e* (**Figure 5D,E**). Relative RNA-seq profiles revealed expression levels of Nt5e/CD73 are significantly elevated in pancreatic ducts with expression of mutant *Kras* compared to wild type ducts (**Figure 5F**). Consistent with previous genetic, proteomic and IHC data showing elevated Ras activity significantly decreases PTEN levels in ductal derived PanIN and loss of PTEN is important in ductal cell transformation(29, 35), we observed a significant decrease in *PTEN* mRNA levels in Tam^10^ ducts compared to Tam^0^ ducts (**Figure 5G**). These data implicate PTEN as an upstream suppressor of CD73. To determine if members of the AKT pathway were elevated in ducts with PTEN loss, we analyzed relative RNA-seq transcriptomic levels of Akt1, Akt2 and Akt1s1 which were all significantly elevated in Tam^10^ ducts (**Figure 5H**). These data indicate elevated CD73 is an early response to oncogenic mutant Kras and K-Ras activity in pancreatic ducts and is associated with loss of PTEN and elevated AKT signaling. In addition to elevated CD73, we observe high expression of *Cd274*/PD-L1, *Arg1*, *Smad7* and *Lgals3*/Galectin3, all mediators of immunosuppression, in *Kras* mutant ducts (**Figure 5I**).

**Figure 5.**
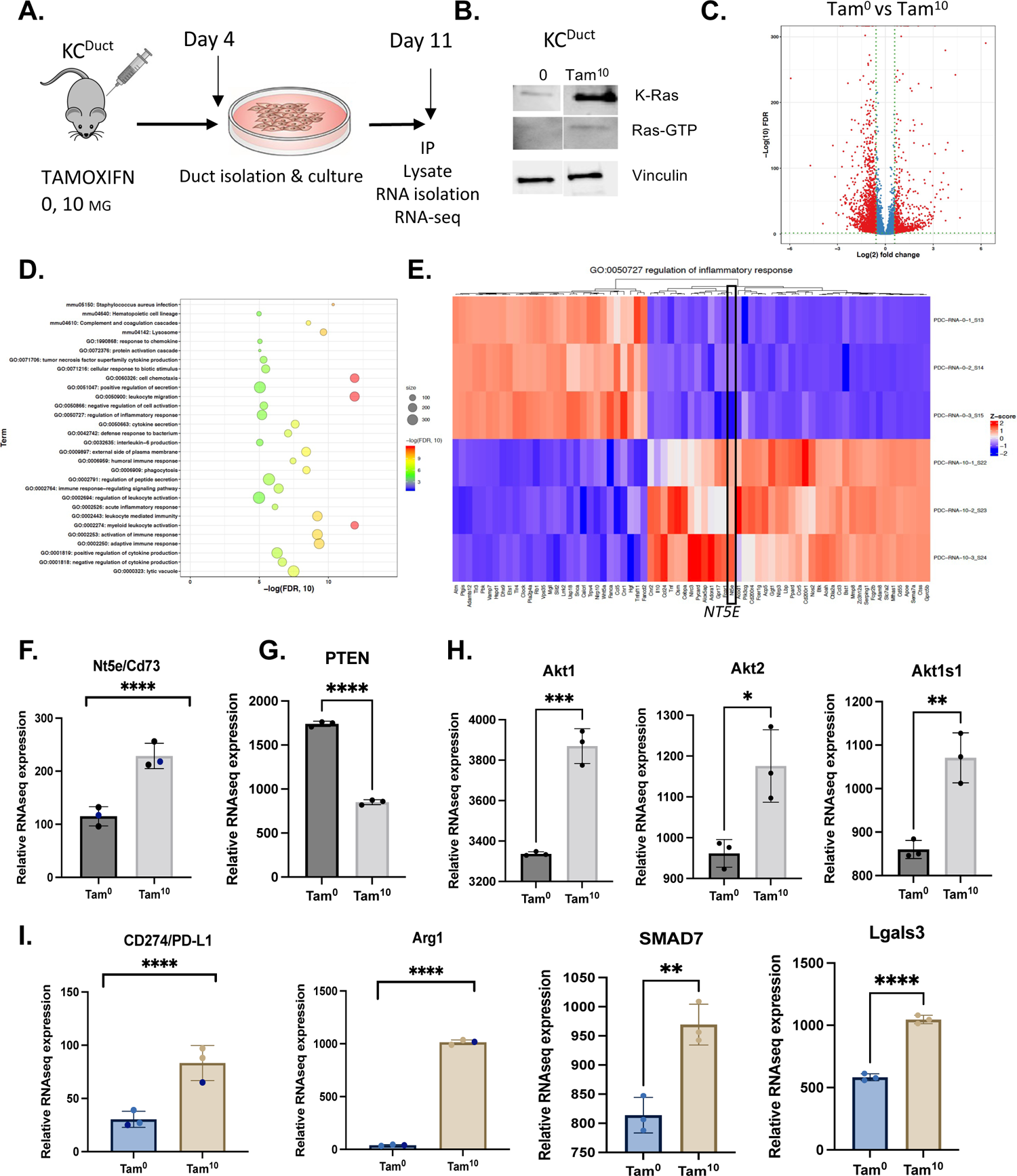
Oncogenic *Kras* expression significantly increases transcriptomic levels of CD73 and significantly reduces transcriptomic levels of PTEN. **A)** Schematic of experimental setup to generate whole transcriptomic profiles of Kras mutant pancreatic ducts. **B)** Western blot showing increased expression of K-Ras in *ex vivo* cultured pancreatic ducts from KC^Duct^ mice. Figure is adapted from Singh et al, 2021. **C)** Violin plot analysis of RNA-seq data generated from Tam^0^ versus Tam^10^ *ex vivo* cultured ducts. **D)** WebGestaltR top 30 pathways enriched in Tam^10^ ducts compared to Tam^0^ ducts. **E)** Gene Ontology Heat Map of highly enriched pathways in Kras mutant ducts. Regulation of inflammatory response is one of the top 30 pathways. *Nt5e* is elevated in regulation of inflammatory response. **F)** Relative RNA-seq signature of Nt5e/CD73 in *ex vivo* cultured pancreatic ducts. CD73 is significantly increased in Kras mutant pancreatic ducts *(****P<0.001*) Student’s t-test (n=3 per group analyzed by RNA-seq). **G)** Relative RNA-seq expression data for PTEN shows a significant reduction in PTEN mRNA levels as a function of K-Ras levels and activity; notably loss of PTEN is associated with elevated CD73. **H)** Akt1, Akt2 and Akt1s1 are all significantly increased in Tam^10^ versus wild type pancreatic ducts. I) *CD274, Arg1, SMAD7 and Lglas3* are significantly elevated in (Tam^10^) *Kras* mutant ducts *(**P<0.01; ****P<0.001*).

### CD73-dependent adenosine levels determine PanIN and PDAC progression

To determine if CD73 activity was important for ductal-derived metaplasia and transformation to PanIN and PDAC, we treated GEM KC^Duct^ and KC^Acinar^ mice with Adenosine 5’-(α,β ethylene) diphosphate (APCP) a small molecule inhibitor of CD73, which was started after Tamoxifen injections (**Figure 6A**) and continued administration every other day for the duration of the experiment. Inhibition of CD73 resulted in a significant reduction in surface occupied by PanINs or PDAC in KC^Duct^ mice, but not KC^Acinar^ pancreata, confirming the functional relevance of CD73 in ductal cell transformation (**Figure 6B**). We used IHC to evaluate Kras downstream pathways in pancreata from vehicle versus APCP treated KC^Duct^ mice and observed reduced staining intensity for p-ERK^202/204^ and p-AKT^T308^ indicating autonomous adenosine signaling drives ductal cell transformation (**Figure 6C**). A significant increase in CD8+ cells infiltration was found in pancreata from APCP treated KC^Duct^ mice compared to vehicle treated mice which indicates reduced adenosine generation permits infiltration of CD8+ T cells and perturbs ductal cell transformation (**Figure 6C,E**).

**Figure 6.**
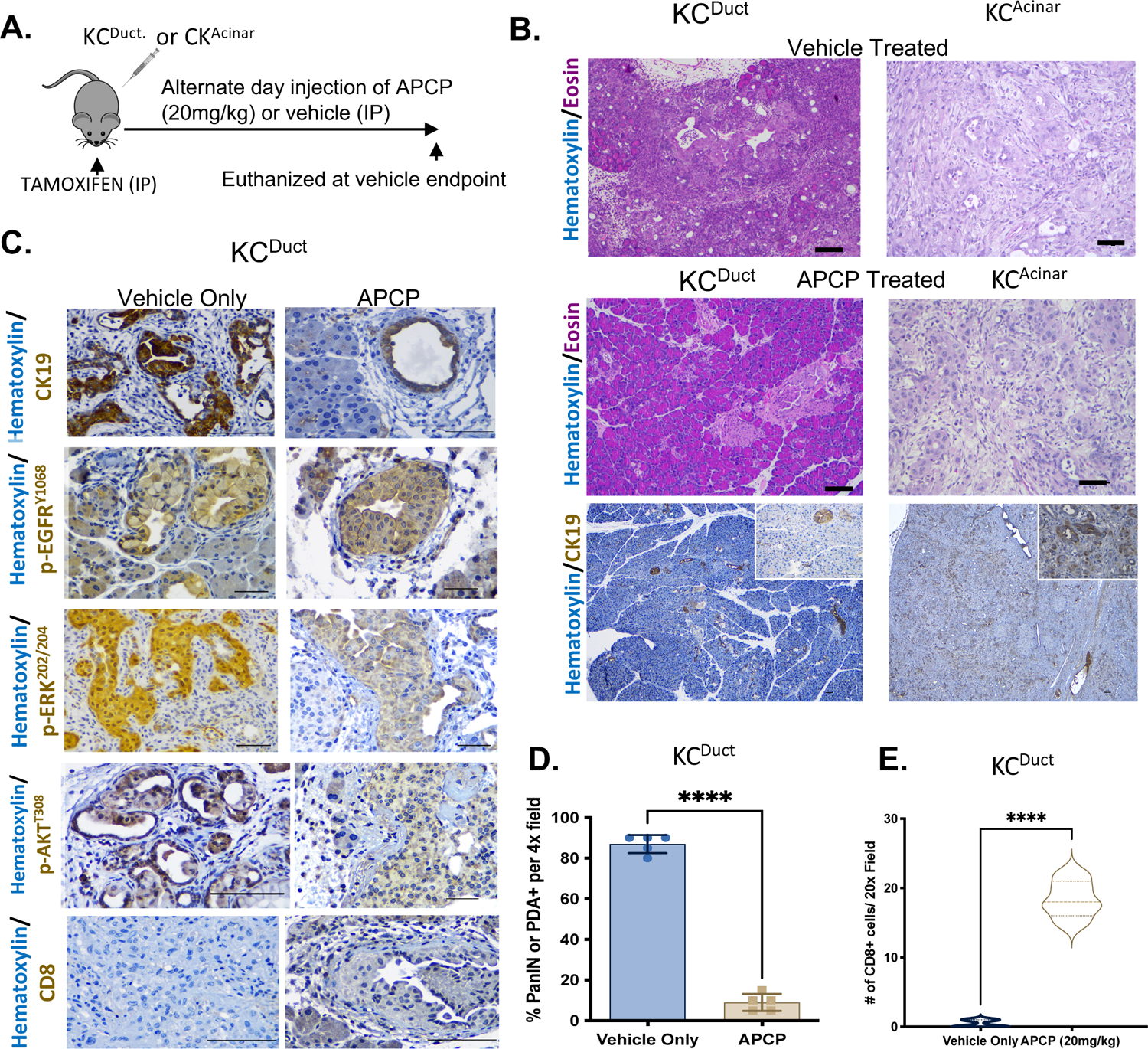
Inhibition of CD73 using intraperitoneal delivery of APCP significantly reduced ductal-derived PDAC. **A**) Schematic of preclinical model to evaluate the requirement for CD73 in a spontaneous GEM model of ductal derived PanIN and PDAC. **B)** Inhibition of CD73 significantly reduced PanIN and PDAC in KC^Duct^ but not KC^Acinar^ GEM mice. **C**) Inhibition of CD73 reduced the IHC staining intenstisty of pERK^202/204^ and pAKT^T308^ in KC^Duct^ pancreata, but did not alter p-EGFR^Y1068^ **D)** Inhibition of CD73 significantly reduced the percentage of pancreatic area positive for PanIN or PDAC (\*\*\**P<0.0001*) and **E)** significantly increased the number of CD8+ cells per 20x field analyzed (*P<0.001*). Scale bars are 100um.

We then aimed to determine if inhibition of CD73 in murine KPC cells generated from *Pdx:Cre; LsL-Kras^G12D^; LsL-Trp53^R173H^* mice *in vitro* would reduce their proliferative capacity. APCP treatment *in vitro* resulted in a significant decrease in KPC cell line proliferation at 72 hours (**Figure 7A**) indicating autonomous adenosine signaling is important for murine PDAC proliferation. To determine if pharmacologic inhibition of adenosine generation using APCP would decrease tumor growth *in vivo*, we performed subcutaneous (subQ) injections of murine KPC cells into syngeneic C57BL/6 mice (**Figure 7B**). IHC analysis confirmed KPC cells express CD73 and KPC tumors have significantly higher levels of adenosine than wild type pancreas (**Figure 7C-D**). Our data revealed peritumor APCP treatment of KPC cells significantly decreases final tumor volume (**Figure 7E**). HPLC analysis was used to confirm APCP significantly reduces intratumoral adenosine levels *in vivo* (**Figure 7F**). Immunohistochemical and flow-cytometry analysis revealed a significant increase in infiltration by granzyme B+ cells (**Figure 7I**), CD45+ and CD8+ T cells in tumors upon APCP treatment vs control (**Figure 7K-L**). In addition, we observed significantly increased CD8αTCRαβ+ cells in the spleen of APCP treated mice (**Figure 7M**).

**Figure 7.**
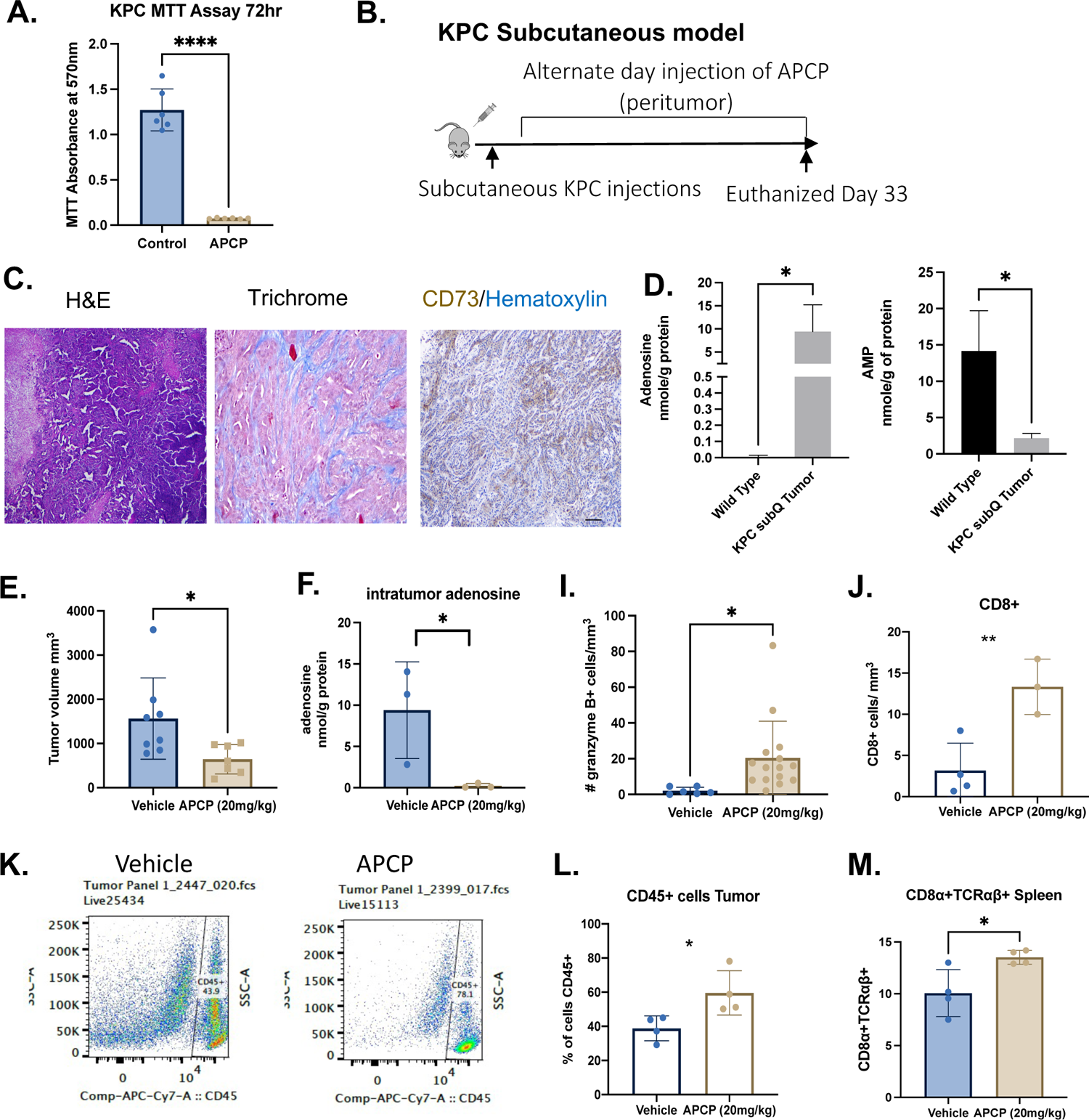
Inhibition of CD73 using peritumor delivery of APCP significantly reduced intratumoral adenosine concentrations and elevated anti-tumor immunity. **A)** MTT assay showing CD73 inhibition significantly reduces KPC proliferation rate *in vitro* (*P<0.001*). **B)** Schematic of KPC subcutaneous tumor injection model. **C)** IHC image of CD73+ KPC tumor and **D)** HPLC analysis of adenosine and AMP levels in KPC subcutaneous tumors compared to wild type pancreata. KPC tumors have significantly increased levels of adenosine and decreased levels of AMP compared to wild type pancreas *(*P<0.05). **E**)* Inhibition of CD73 significantly reduces final tumor growth volume in the KPC subcutaneous model (*\*P<0.05*). **H)** Inhibition of CD73 significantly reduces intratumoral levels of adenosine *(*P<0.05*) and significantly increases the number of (I) granzyme B+ cells and intratumoral **(J)** CD8+ T cells. **K)** We used FACS to analyze the percentage of CD45+ cells and observed a significant increase in CD45+ cells in tumors from APCP treated mice compared to vehicle treated mice **(L)** (P<0.05) and **(M)** a significant increase in splenic CD8a+TCRaB+ cells (\**P<0.05*). Statistical analysis were performed using a student’s unpaired t-test in Prism GraphPad software.

### CD73 dependent generation of adenosine is a potent mediator of immunosuppression in PDAC

In addition to APCP, we tested AB680, another CD73 small molecule inhibitor. Clinical trials using AB680 from Arcus Bioscience have recently shown AB680 in combination with NP/Gem plus zimberelimab had an overall response rate of 41%(45). Using the KPC subcutaneous model, we injected 200K KPC cells subcutaneously, then initiated oral gavage delivery of AB680 three days per week until mice needed to be euthanized (**Figure 8A**). At the conclusion of the experiment, we quantified a significant reduction in tumor volume in AB680 treated mice compared to vehicle treated alone (**Figure 8B**). To directly assess individual tumor responses, we graphed tumor volumes per week which revealed AB680 treatment significantly increased tumor doubling time (**Supplemental Figure 5**). HPLC analysis confirmed a significant decrease in intratumoral adenosine levels in AB680 treated mice indicating oral gavage delivery method successfully reduces CD73 activity *in vivo* (**Figure 7C**). To determine if immunosuppression was altered in AB680 treated KPC tumors, we performed cytometry by time of flight (CyTOF) immunoprofiling of the tumors at time of euthanasia. CyTOF analysis revealed a significant increase in activated CD8 T cells, activated CD4 T cells, dendritic cells, macrophages and myeloid derived suppressor cells (MDSC) (**Figure 8D-I**). Analysis of immune checkpoint markers indicated activated CD4+ and CD8+ T cells increased expression of PD-1 in AB680 treated mice (**Figure 8G**). We used IHC to stain for granzyme B which confirmed an increase in granzyme B+ cells in tumors from AB680 treated mice (**Supplemental Figure 5**). Consistent with data from human clinical trials, we did not observe elevated liver toxicity in AB680 treated mice (**Supplemental Figure 5**).

**Figure 8.**
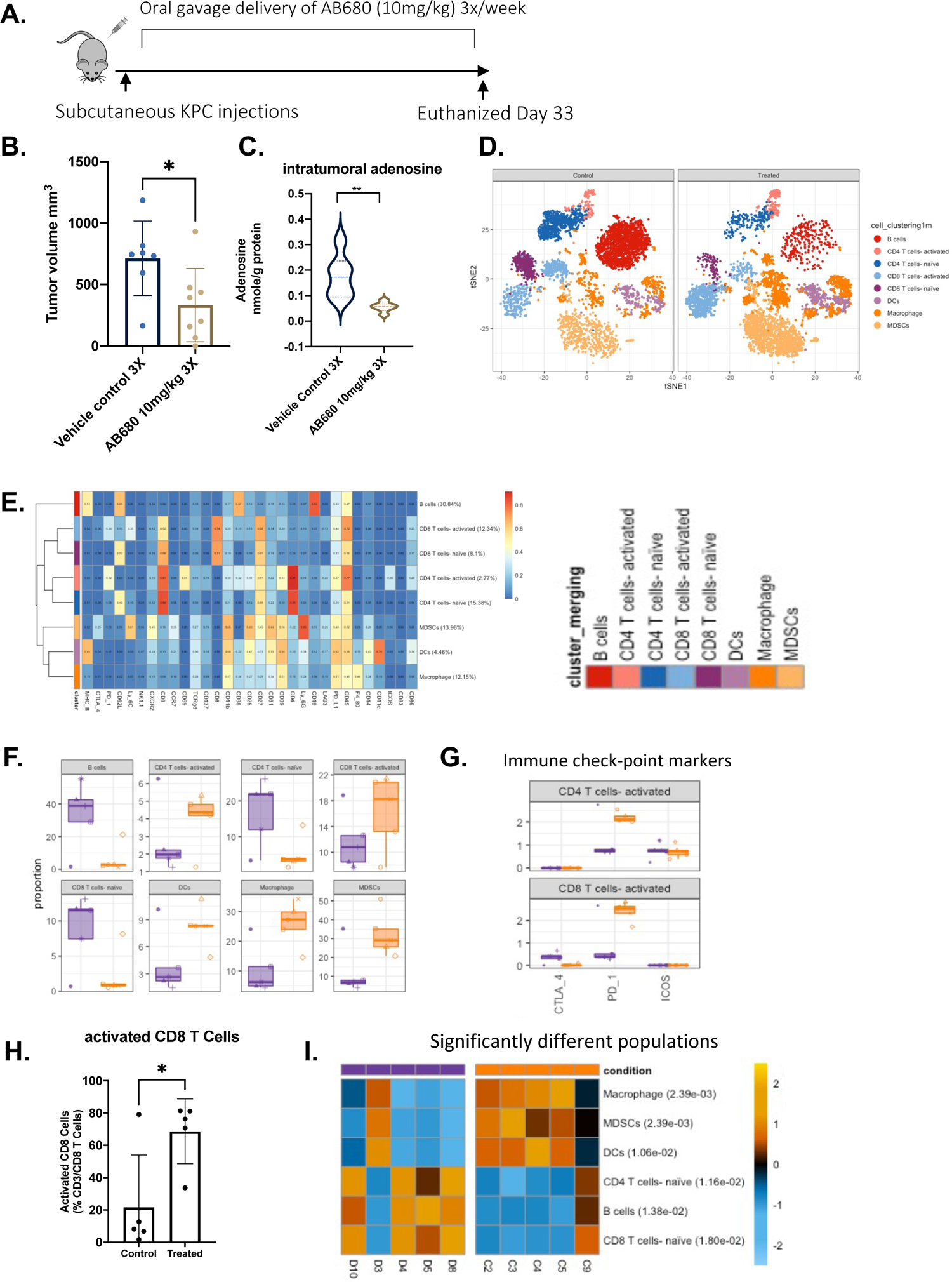
AB680 oral gavage treatment reduces tumor KPC tumor growth rate and elevates intratumoral activated CD8+ T cells. **A)** KPC subcutaneous tumors were analyzed weekly (n=10 per group). **B)** At the conclusion of this experiment, AB680-treated mice had significantly smaller tumor volume than vehicle control-treated mice. *\*P<0.05*. Statistical analysis was performed using a student’s t-test in Prism Graphpad software **C)** HPLC analysis shows a significant decrease in adenosine levels in tumors from AB680 treated mice versus vehicle treated mice. *\*\*P<0.01* compared to vehicle controls. (n=8 per group). **D**) CyTOf vSNE plots by group show elevated clusters of activated CD8 T cells, CD4 T cells, MDSCs and Macrophages. **E**) Heat Map analysis of CyTOF data showing relative expression of cell clusters. **F**) Quantitative global population analysis reveal increased activated CD4 T cells, CD8 T cells, macrophages and MDSCs. Notably, activated CD4 and CD8 T cells increased expression of PD_1. **H**) Quantitative analysis of significantly increased activated CD8 T cells and **I)** Heat map analysis showing significant increases in macrophage, MDSC, DCs in tumors from mice treated with AB680, indicating increased anti-tumor immunity in AB680-treated mice compared to control mice. *\*P<0.05* and a student’s test using Prism Graphpad software was used to calculate statistics.

## METHODS

### RNA preparation and sequencing

Samples were rinsed in PBS and immediately frozen in liquid nitrogen. Total RNA was extracted using a miRNeasy Mini Kit (74104, Qiagen) and submitted to the Cancer Genomics Center at the University of Texas Health Science Center. Total RNA quality was measured using Agilent RNA 6000 Pico kit (#5067-1513) by Agilent Bioanalyzer 2100 (Agilent Technologies, Santa Clara, USA). The samples with RNA integrity number (RIN) greater than 7 were used for library preparation. Libraries were prepared following the manufacturer’s instructions of the Roche KAPA mRNA HyperPrep Kit (KK8581) and the KAPA Unique Dual-indexed Adapter Kit (KK8727). The quality of the final libraries was examined using Agilent High Sensitive DNA Kit (#5067-4626) by Agilent Bioanalyzer 2100 (Agilent Technologies, Santa Clara, USA), and the library concentrations were determined by qPCR using Collibri Library Quantification kit (#A38524500, Thermo Fisher Scientific). The pooled libraries were sequenced on the Illumina NextSeq 550 platform using the pair-ended 75 bp by a 150-cycle High Output v2.5 Kit (#20024907, Illumina, Inc., USA). We used ultrafast universal RNA-seq aligner STAR (v2.5.3a) to map the RNA-seq reads to mouse reference genome GRCm38(46). To obtain the uniquely-mapped reads per gene from the GencodeM15 (GRCm38) reference, we set the argument –quantMode to “GeneCounts”. We filtered out those genes with < 5 reads in all samples and conducted the differential expression analysis for the remaining genes by DESeq2 software(47). The p-values of genes were adjusted using the Benjamini and Hochberg’s procedure to control the false discovery rate (FDR). And the differentially expressed genes were defined as the genes with absolute log2 (fold change) > 0.58 and FDR < 0.05. None redundant Gene Ontology (GO) and KEGG pathway enrichment analysis were performed using WebGestalt (v0.4.3) software(48). All raw data and processed read count have been submitted to Gene Expression Omnibus (GSE189130).

### Basal and Acinar signature genes derived from mouse in human homologs

We adapted the differential expressed genes analysis from our recent cell of origin GEM models between Ductal tumor and Acinar tumor in mouse(27). We defined those genes (304) with log2 (Fold change) > 2 and FDR < 0.01 as the signature of Ductal signature genes (917). We further defined Acinar signature genes as those genes with log2 (Fold change) < −2 and FDR < 0.01. We then downloaded the latest Mouse Human homolog gene symbol (v7.4) file from https://data.broadinstitute.org/gsea-msigdb/msigdb/annotations_versioned/ (access by 4/20/2021). We obtained the mouse-derived human homologs signatures for Ductal (271) and Acinar tumor genes (877), respectively.

### Gene Set Variation Analysis (GSVA)

We obtained RNA-seq datasets and differentially expressed genes signatures from two independent PDACC patient cohorts, primary PDACCs of high cellularity from the Australian International Cancer Genome Initiative (ICGC) (access by 4/21/2021) and primary PDACCs from The Cancer Genome Atlas (TCGA) Research Network (access through TCGA data portal by 4/22/2021) (16, 17). Specifically, immunogenic, squamous, classical and basal subtypes classification were considered as previously described by TCGA (Supplementary table S1 in (16), GUID: C0853F1D-79F0-4125-BA86-8545A54FA572). We used the construct the signature genes using the aforementioned human homolog Ductal and Acinar tumor gene sets. Then, we conducted the GSVA analysis with default setting for immunogenic and squamous samples and basal and classical samples, respectively(49). The GSVA score indicates the relative variation of signature genes activity over the AU and TCGA samples.

### Venn diagram analysis

Differentially expressed genes between Immunogenic vs. Squamous and Classical vs. Basal subtypes were obtained from (17) (Suppl. Table 17) and TCGA (16) (Suppl. S1 table, GUID: 3DD81EAF-3FD4-48CE-A9DA-2454820DAB10). Upregulated genes in human Squamous and Basal subtypes were compared to upregulated genes in KPC^Duct^ mouse model and shown as Venn diagram analysis.

### Imaging mass cytometry (IMC)

Metal-labeled antibodies were prepared according to the Fluidigm protocol. Antibodies were obtained in carrier/protein-free buffer and then prepared using the MaxPar antibody conjugation kit (Fluidigm). After determining the percent yield by absorbance measurement at 280 nm, the metal-labeled antibodies were diluted in Candor PBS Antibody Stabilization solution (Candor Bioscience) for long-term storage at 4°C. Antibodies used in this study are listed in Table E1 (50).

Tumor sections were baked at 60°C overnight, then dewaxed in xylene and rehydrated in a graded series of alcohol (ethanol absolute, ethanol:deionized water 90:10, 80:20, 70:30, 50:50, 0:100; 10 minutes each) for imaging mass cytometry. Heat-induced epitope retrieval was conducted in a water bath at 95°C in Tris buffer with Tween 20 at pH 9 for 20 minutes. After immediate cooling for 20 minutes, the sections were blocked with 3% bovine serum albumin in tris-buffered saline (TBS) for 1 hour. For staining, the sections were incubated overnight at 4°C with an antibody master mix. Samples were then washed 4 times with TBS/0.1% Tween20. For nuclear staining, the sections were stained with Cell-ID Intercalator (Fluidigm) for 5 minutes and washed twice with TBS/0.1% Tween20. Slides were air-dried and stored at 4°C for ablation.

The sections were ablated with Hyperion (Fluidigm) for data acquisition. Imaging mass cytometry data were segmented by ilastik and CellProfiler. Histology topography cytometry analysis toolbox (HistoCAT) and R scripts were used to quantify cell number, generate tSNE plots, and perform neighborhood analysis. 40 For all samples, tumor and cellular densities were averaged across 3 images per tumor, with n = 3 per group.

### Mass cytometry (CyTOF)

Tumor tissues were harvested and digested with 1 mg/ml collagenase P (MilliporeSigma) and 0.5 mg/ml DNase I (MilliporeSigma). Single-cell suspensions were stained with 5 μM Cell-ID Cisplatin (Fluidigm Corp.) and incubated with Fc block (BD Biosciences), followed by surface antibody cocktail. Antibody details including final concentrations can be found in **Supplemental Table 3**. Next, cells were washed and fixed in Maxpar Fix I buffer (Fluidigm Corp.) and barcoded using the Cell-ID 20-Plex Pd Barcoding Kit (Fluidigm Corp.). Next, the cells stained with 1.25 μM Cell-ID Intercalator-Ir (Fluidigm Corp.) overnight. Sample acquisition was performed on a Helios mass cytometer (Fluidigm Corp.). The analysis was performed using R package CyTOF Workflow (51) and FlowJo version 10 software (FlowJo LLC).

### Cell lines

HPNE, Capan-2 and ASPC1 cells were purchased from ATCC and were maintained following vendor’s instructions. Murine pancreatic adenocarcinoma cells (KPC) were a generous gift from Dave Tuveson (Cold Spring Harbor Laboratory, Cold Spring Harbor, NY) and were maintained in DMEM (Thermofisher Scientific-10567014) with 10% Fetal bovine serum. All cell lines were maintained at 37^0^C and 5% CO_2_ in a humified incubator.

### Animal Models

All animal experiments were approved and performed under the guidelines of The Center for Laboratory Animal Medicine and Care (CLAMC) at University of Texas Health Science Center at Houston. C57BL/6 (000664) mice were purchased from Jackson laboratories. Hnf1b;Cre^ERTM^ and Ptf1a;Cre^ERTM^ mice were purchased form Jackson Laboratories. KPC^Duct^ and KPC^Acinar^ mice were generated by crossing to LSL-Kras^G12D^ and LSL-TP53^R172H^ as described previously (27). For obtaining ectopic Kras expression, transgenic mice with CAG-lox-GFP-stop-lox-KrasG12V (52) were received from Craig Logsdon, MD Anderson Cancer Center, Houston, TX. Strains of Hnf1b;Cre^ERTM^ mice were crossed with cLGL-KRAS^G12V^ to generate cK^Duct^ mice and obtain mutant cKras expression in adult pancreatic ductal cells. Similarly, Ptf1a;Cre^ERTM^ were crossed with cLGL-KRAS^G12V^ mice to generate cK^Acinar^ mice to obtain cKras expression in mature acinar cells. These mice express GFP in whole body and lose GFP after Cre mediated recombination. Mice were genotyped by PCR or Transnetyx. To achieve different levels of mutant cKras in ductal cells, mice at an age of 6-8 weeks were injected with 1mg tamoxifen (Sigma, T5648) subcutaneously one day (1mg dose), 5mg tamoxifen for 1 day (5mg dose) and 5mg tamoxifen for 2 consecutive days (10 mg dose). These mice were named as Tam^1^, Tam^5^ and Tam^10^ respectively. An n=10-12 mice were evaluated for each tamoxifen dose. For subcutaneous xenograft models, 1X10^5^ KPC cells in PBS: Matrigel mix (1:1) were injected in the left flank of C57BL/6mice. Tumor size was calculated twice a week with vernier caliper. Tumor volume was calculated as length × width × width/2 in cubic millimeters. Tumor doubling time was calculated using the method described previously(53).

### Histopathology

Formalin fixed and sectioned pancreas tissue was deparaffinized with histoclear followed by hydration with ethanol and water and staining with hematoxylin. Sections were next counterstained with Eosin and dehydrated stepwise with ethanol and histoclear. Slides were mounted with coverslip using a mounting medium. All pancreatic pathologies in the genetic engineered models were classified by pathologists at University of Texas Health Science Center.

### CD73 inhibitor administration

APCP ( αβ-methylene ADP, Sigma-Aldrich; catalog no. M3763) was purchased from Sigma. AB680 was purchased from MedChemExpress (catalog no. HY-125286). Mice bearing subcutaneous KPC tumors or mice with spontaneous PDAC tumors were treated with CD73 inhibitors. 20mg/kg APCP in PBS or vehicle control (PBS) was administered IP for spontaneous model and peri-tumor in subcutaneously model. Mice were given APCP on alternate days until the end of experiment. For AB680 treatment, the stocks were prepared in 100% DMSO. For oral gavage AB680 was diluted in 10%DMSO+90% SBE beta cyclodextrin (SBE-b-CD) in 0.9% saline. 10mg/kg AB680 or vehicle control (10%DMSO+90% SBE-b-CD in 0.9% saline) were given by oral gavage on alternate days until the end of experiment.

### Ras activity pull down assay

Ras activity in the pancreas tissue lysates was performed using the Active Ras pull down and detection kit (Thermofisher Scientific). Up to 30 mg of tissue (fresh or frozen at −80°C) was washed with 1X cold PBS, was homogenized in 1 ml lysis buffer (Cell Signaling Technology, 9803S) containing protease inhibitor cocktail (Roche, 4693159001). Lysates were placed on ice for 30 minutes followed by sonication for 2 minutes with 10 seconds on/off cycle and centrifugation for 10 minutes at 10000g at 4°C in a microcentrifuge. The pellet was discarded, and the lysate was used for protein estimation by BCA method. For pulldown, a 500 μg protein equivalent of lysates were incubated with beads coated with Raf1-RBD provided with the kit, for 1 hour at 4°C. Beads were then washed 3 times with ice-cold lysis buffer, and bound protein was eluted for 15 minutes with Laemmli sample buffer that had been preheated to 95°C. Aliquots of lysates were also saved for further quantification of total Ras or protein loading controls by immunoblotting. Pulldown proteins were analyzed by immunoblotting with Ras antibody provided with the kit.

### Western Blot Analysis

Cell and tissue extracts were prepared using cell lysis buffer (Cell signaling #9803S) with protease inhibitor cocktail tablets (Cell signaling #5871). BCA method (Thermofischer Scientific) was used for protein quantification. A 20μg of protein was separated using SDS-PAGE (Bio-Rad). Trans-blot Turbo Transfer kit (Bio-Rad) was used for semi-dry transfer. After transfer, membranes were blocked using 5% skimmed milk (Bio-Rad) in TBST (TBS buffer containing 0.5% Tween-20) for 1 hour followed by overnight incubation with primary antibodies (in 5% milk and dilution as per manufacturer’s instructions) at 4°C. On day-2, the membrane was washed 4 times with TBST buffer and incubated with the respective HRP-conjugated secondary antibody (1:5000) for 1 hour. Further, membranes were washed four times with TBST buffer and developed using ClarityTM Western ECL Substrate (Bio-Rad #1705061). Primary antibodies used in this study are described in **Supplemental Table 4**.

### Primary pancreatic duct culture

Pancreatic ducts were cultured as defined previously (54) from Tam^0^ (WT), Tam^1^, Tam^5^ and Tam^10^ mice four days post-tamoxifen administration. Briefly, Pancreas was collected, minced to 1mm pieces, and digested for 30 min at 37°C in digestive solution (0.1% soybean trypsin inhibitor and 0.1% Collagenase). Cells were filtered through 40μm filter, washed additional two times with culture medium and plated on collagen coated plates in complete medium (DMEM/F12 (Life Technologies 11330-032) 500 mL, Penicillin-streptomycin (100×; Life Technologies 15140-122) 5 mL, 1×Nu-serum IV (BD Biosciences 355104) 25 mL, 5% Bovine pituitary extract (3 mg/mL; BD Biosciences 354123) 4.2 mL, 25 μg/mL ITS+ Premix (BD Biosciences 354352) 2.5 mL, Epidermal growth factor (100 μg/mL; BD Biosciences 354001) 100 μL, 20 ng/mL Cholera toxin (1 mg/mL; Sigma-Aldrich C8052) 50 μL, 100 ng/mL3,3,5-Triiodo-L-thyronine (50 μM; Sigma-Aldrich T2877)50 μL, 5 nM Dexamethasone (100 mM; Sigma-AldrichD1756) 5 μL, 1 μM D-Glucose (Sigma-Aldrich G5400) 2.5 g 4.7 mg/mL, Nicotinamide (Sigma-Aldrich N3376) 0.66 g 1.22 mg/mL and Soybean trypsin inhibitor (type I; Sigma-Aldrich T6522) 50 mg 0.1 mg/mL. The cultures grew to confluency in one week and fibroblast contamination was reduced using differential trypsinization method. Cell lysates were collected, and equal amounts of protein were subjected to Western analysis.

### Immunohistochemistry

Paraformaldehyde fixed and sectioned tissue were baked at 60°C for 30 minutes. Deparaffinization and rehydration was performed using histoclear followed by subsequent incubation with 100%, 70%, 30% ethanol and deionized water. Sections were permeabilized using PBST (PBS containing 0.25% Tween-20) for 10 min and endogenous peroxidases were blocked with 0.3% H2O2 in PBS for 15 min. Antigen retrieval was performed by antigen unmasking solution (Vector Laboratories, H-3300) using heat-mediated microwave method. Sections were blocked with 10% FBS in PBST followed by incubation with primary antibodies overnight at 4°C (Primary antibodies used in this study are described in Supplemental Table 1). Next day, sections were washed three times with PBST and then incubated with secondary antibodies (1:500) at room Temperature for 2 hours. The sections were washed three times with PBST and detection was performed using Vectastain Elite ABC kit (Vector Laboratories, PK-6100) and DAB Peroxidase (HRP) Substrate kit (Vector Laboratories, SK-4100). Sections were counterstained with hematoxylin, mounted with coverslip using mounting media and visualized under light microscopy.

### Nucleoside/Nucleotide extraction and quantification

Cell culture supernatants, mouse serum or tumor tissues were collected at the end of the study. Serum used for adenosine analysis was frozen with adenosine inhibitor cocktail containing 10 μM APCP (CD73 inhibitor; αβ ethylene ADP, Sigma-Aldrich; catalog no. M3763) 10 μM dipyridamole (equilibrative nucleoside transporter inhibitor; Sigma-Aldrich; catalog no. D9766) and 10 μ deoxycoformycin (adenosine deaminase inhibitor; R&D Systems; catalog no. 2033) to preserve nucleosides.

Tumors were flash frozen with liquid nitrogen and stored at −80 °C. Cell supernatants, serum or tumor protein lysate was extracted with perchloric acid and neutralized with KHCO_3_/KOH. Samples were acidified with ammonium dihydrogen phosphate and phosphoric acid. Reaction supernatant was collected by centrifugation. Extracted samples were analyzed by reversed phase high performance liquid chromatography (RP-HPLC)(55). Representative AMP and adenosine peaks were identified and measured using the respective standard HPLC curve. For tumors adenosine levels were normalized to the protein levels in the tumor lysate. To determine if PDAC cell lines had increased CD73 activity, cell line supernatant was replaced with HBSS and 250ng AMP were added. To evaluate CD73 activity, we extracted 100ul of HBSS supernatant at time 0,10, 30, 60 and 120 minutes.

### RNA isolation and quantitative RT-PCR

Total RNA was extracted with the RNeasy RNA isolation kit (Qiagen) and reverse transcribed with a cDNA Reverse Transcription Kit (Bio-Rad). Quantitative RT-PCR was performed with SYBR Green Master Mix (Bio-Rad) on a Bio-Rad real-time PCR system. PCR primer sequences used in the study were obtained from PrimerBank (https://pga.mgh.harvard.edu/primerbank/) and were synthesized at Integrated DNA Technologies. GAPDH was used as housekeeping gene and the expression levels of mRNA of interest were normalized to GAPDH.

### TCGA Analysis

The Kaplan-Meier curves were generated using TCGA RNA-sequencing data (FPKM_UQ) for PDACC samples, after excluding PNETs and non-PDACC samples. Higher and lower expression levels were stratified on the basis of average expression. Statistical analysis was performed using log-rank tests, and HRs were calculated using Prism software (GraphPad Software, Inc.).

## DISCUSSION

In this study, we used a combination of genetically engineered and syngeneic preclinical models to evaluate the role of adenosine generation in pancreatic cancer. Initially, to define therapeutic vulnerabilities in PDAC based on cell of origin and PDAC subtypes, we used RNA-seq to generate whole transcriptomic profiles of acinar and ductal derived PDAC in mice. Using GSVA scores, we determined that murine tumors arising in pancreatic ducts have significantly overlapping gene expression profiles compared to published human PDAC Squamous and Basal subtypes. Very early in these studies, we also identified *Nt5e*/CD73 was expressed at significantly higher levels in our KPC^Duct^ compared to KPC^Acinar^ PDAC mouse models. This led us to hypothesize neoplastic pancreatic ducts increase extracellular levels of adenosine, an important anti-inflammatory nucleoside involved in immunosuppression and transformation.

In the normal pancreas, divergent data has been published related to expression of ectonucleotidases in ductal epithelium. Analyses of pancreatic juice have identified soluble CD73 and implicate pancreatic ducts express CD73 and CD39 and are critical regulators of adenosine and purinergic signaling in pancreatic homeostasis (56). Studies relying on immunohistochemistry have reported normal pancreas does not stain positive for CD73 indicating in physiologic conditions, CD73 levels are low in exocrine pancreatic cells (57–59). In contrast, in human PDAC specimens, CD73 is highly expressed in a subset of patients and divergent patterns of immunolabeling have been reported(57, 59, 60). Immunolabeling has shown in some patients, CD73 levels are high in malignant epithelial cells with limited staining in adjacent stroma. In other contexts, CD73 expression is observed in vascular endothelium, fibroblasts and infiltrating immune cells(59, 60). In our GEM models, we observe high immunolabeling for CD73 in the luminal side of PanIN and PDAC arising in ductal cells, but not in metaplastic acinar cells, consistent with the observation that acinar derived PDAC has a more immunogenic signature in part due to a lower CD73 RNA-seq profile. In sections from KPC^Acinar^ pancreata, we do observe infiltration of immune cells that stain positive for CD73. These findings are significant as subsets of T lymphocytes are known to express both CD39 and CD73 and can modulate local and systemic concentrations of ATP and adenosine and may alter responses to immunotherapy(61).

Our studies are timely as decades of research have shown extracellular ATP is predominately proinflammatory while the extracellular nucleoside metabolite adenosine is anti-inflammatory(8, 62–64). Integration of purinergic and adenosine signaling is thus a delicate balance to promote tissue homeostasis and repair. Emerging data in cancer biology has shown paracrine purinergic and adenosine signaling mediates immunosuppression in a number of malignancies and has promising therapeutic implications(7, 65–73); yet the role of extracellular adenosine signaling in the initiation of pancreatic cancer has not been studied. Surprisingly, we observe pancreatic ducts elevate CD73 in response to *Kras* mutations, elevated K-Ras signaling and increased CD73 is associated with loss of PTEN. We employed *in vivo* and *in vitro* methods to start undercovering the implications of elevated epithelial expression of in pancreatic ductal transformation. We state adenosine generation is a key distinctive feature of ductal transformed epithelium and is important for PDAC cell survival as well as augmenting immunosuppression. Activation of immune cells after CD73 inhibition, concomitant with reduced spontaneous PanIN lesions arising in pancreatic ducts, highlight cell autonomous and non-cell autonomous mechanisms of extracellular adenosine signaling are critical to the development of ductal pancreatic cancer and initiate early in the setting of the disease. These data further support the idea that elevated CD73 acts as a master regulator in shaping the immunosuppressive PDAC subtypes. In contrast, in tissue from neoplastic acini, we observe elevated AMP, suggesting in acinar cell transformation, AMP is generated by conversion of extracellular ATP to AMP; however, in the absence of sufficient CD73, chronic inflammation and purinergic signaling promote an immunogenic subtype which may dictate better prognosis in human PDAC.

In this work we show for the first time, both *in vitro* and *in vivo*, that increased ductal CD73-dependent adenosine generation is a key driver in the initiation of duct cell malignant transformation and PDAC-associated immune suppression. Hence, our data strongly support that targeting CD73/Adenosine pathway successfully prevents ductal cell injury and improves both antitumor immunity and tumor progression. Further characterization of CD73 and adenosine levels in patients with subtypes of pancreatic cystic lesions, chronic pancreatitis and other etiologies of pancreatic inflammation will augment these studies to determine if patients with these inflammatory conditions which can elevate risk of developing pancreatic cancer may benefit from targeting purinergic or adenosine signaling.

## AUTHOR CONTRIBUTIONS

Data Curation: K.S., E.Y.F., Y.D., V.C., E.V., T.M., M.P., T.C. Resources: L.V., M.I.S., S. S., A. M., P.H.B., H.K.E., D.B.S., Z.Z., J.S.B., G.B. Writing-reviewing and editing: N.C.T., C.J.W., H.K.E., K.L.P. Conceptualization: F.M., J.B.L. Supervision, Funding acquisition: J.B.L. and F.M. Writing original draft: J.B.L.

