## Supplemental Figures for "Pancreatic cancer ductal cell of origin drives CD73-dependent generation of immunosuppressive adenosine"

**A.**

**IPA Top Disease and Function Analysis**

*Hereditary disorder, Ophthalmic Disease, Organismal Injury and Abnormalities*

**B.**

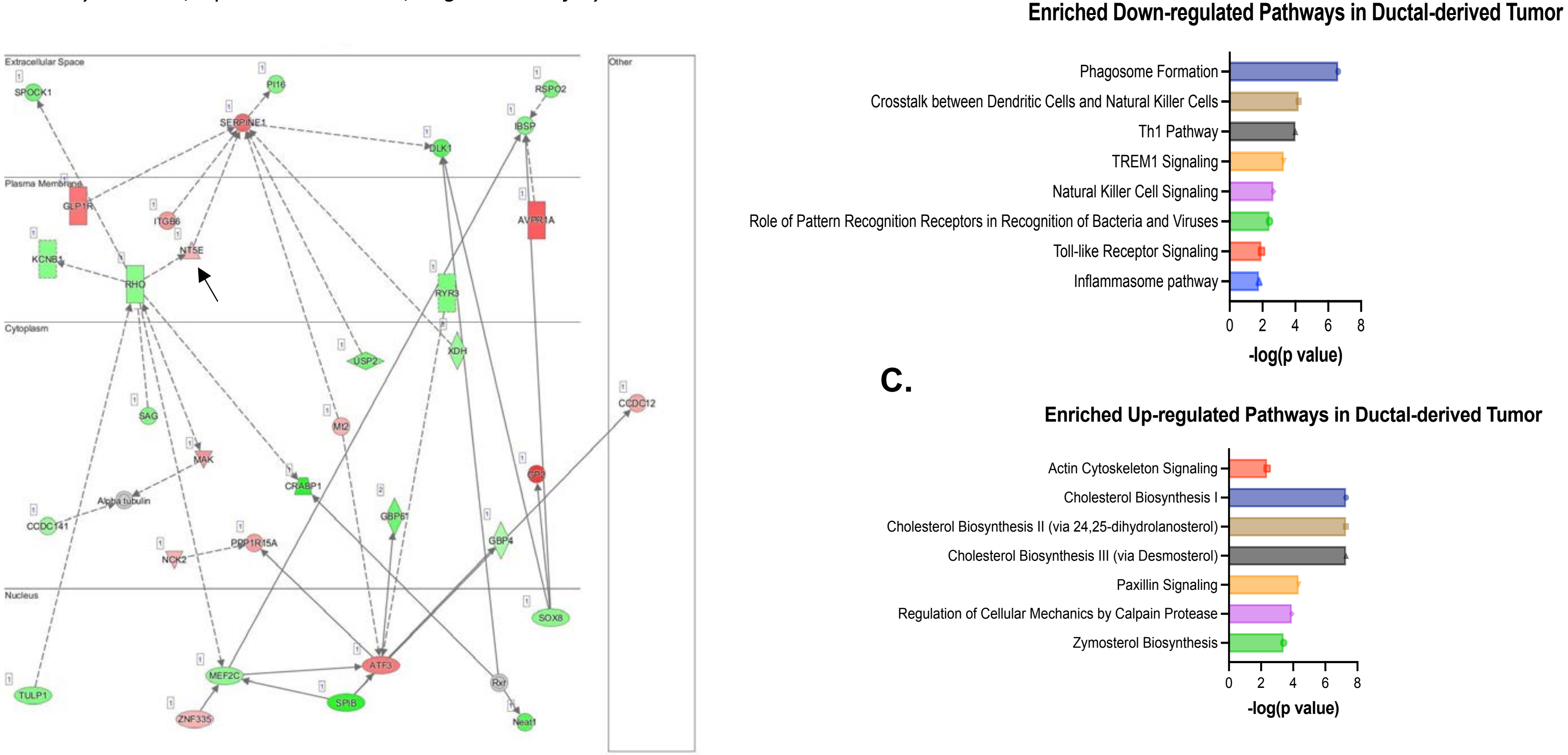


**Supplemental Figure 1. IPA analysis of ductal versus acinar derived PDAC tumors. A)** IPA comparative analysis of murine $KPC^{Duct}$ and $KPC^{Acinar}$ RNA-seq signatures. IPA diagram of Hereditary disorder, Ophthalmic Disease, Organismal Injury and Abnormalities, a Top Disease and Function pathway elevated in murine ductal derived tumors. NT5E (arrow) is significantly elevated in this pathway analysis. **B)** Analysis of whole transcriptomic profiles revealed significant increases in pathways related to cholesterol biosynthesis and **C)** significant decreases in pathways related to innate immunity including Crosstalk between dendritic cells and Th1 Pathway in $KPC^{Duct}$ tumors compared to $KPC^{Acinar}$ tumors.

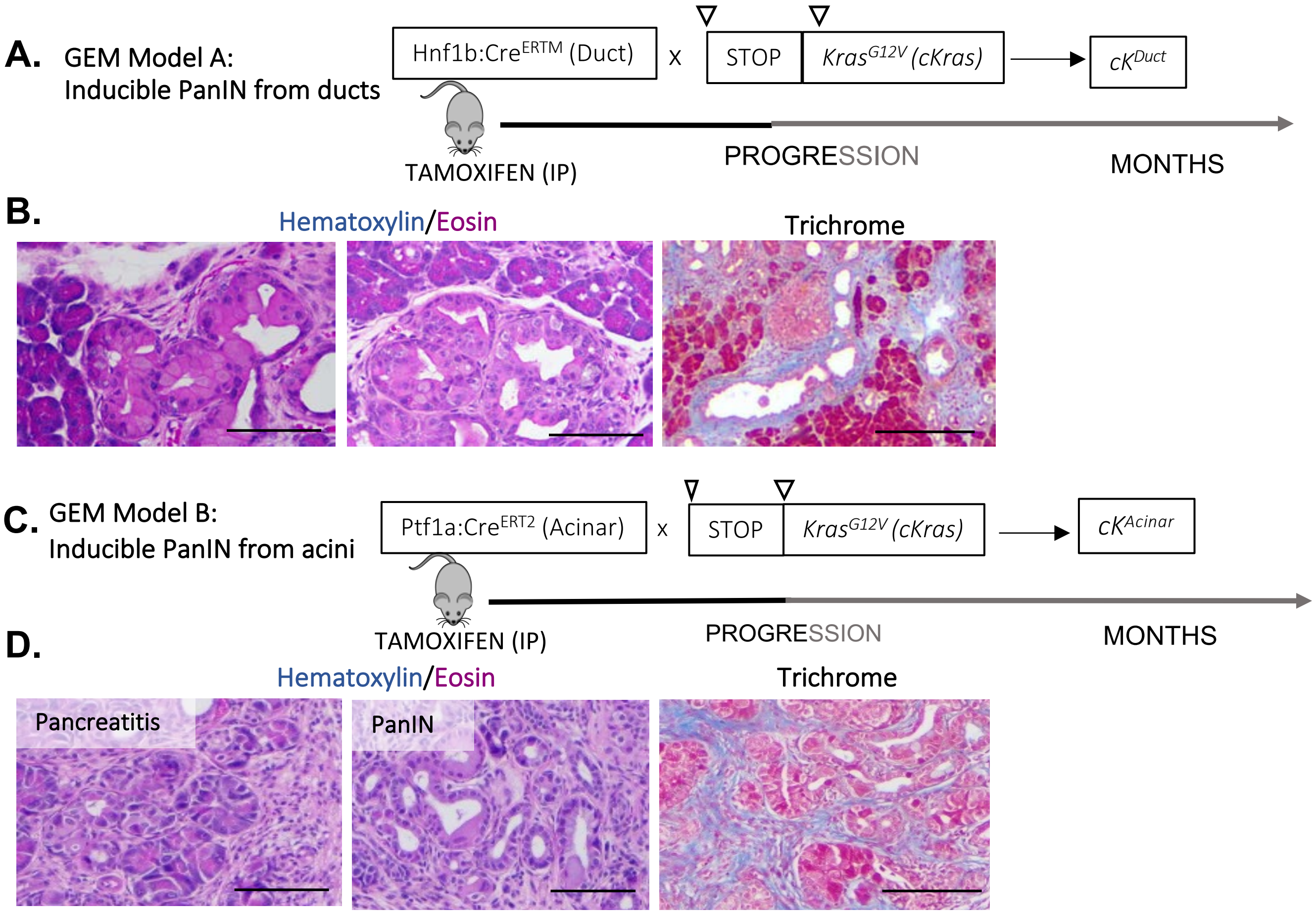


**Supplemental Figure 2. Histologic analysis of PanIN and stroma in Kras mutant expressing ducts and acini. A)** Experimental design to study intraductal PanIN and PDAC. **B**) H&E and Trichrome staining show early and advanced PanIN and collagen deposition adjacent to intraductal PanIN. **C)** Experimental design to study PanIN and PDAC arising in acini. **D)** H&E and Trichrome staining show early and advanced PanIN and collagen deposition adjacent to PanIN arising in acinar cells.

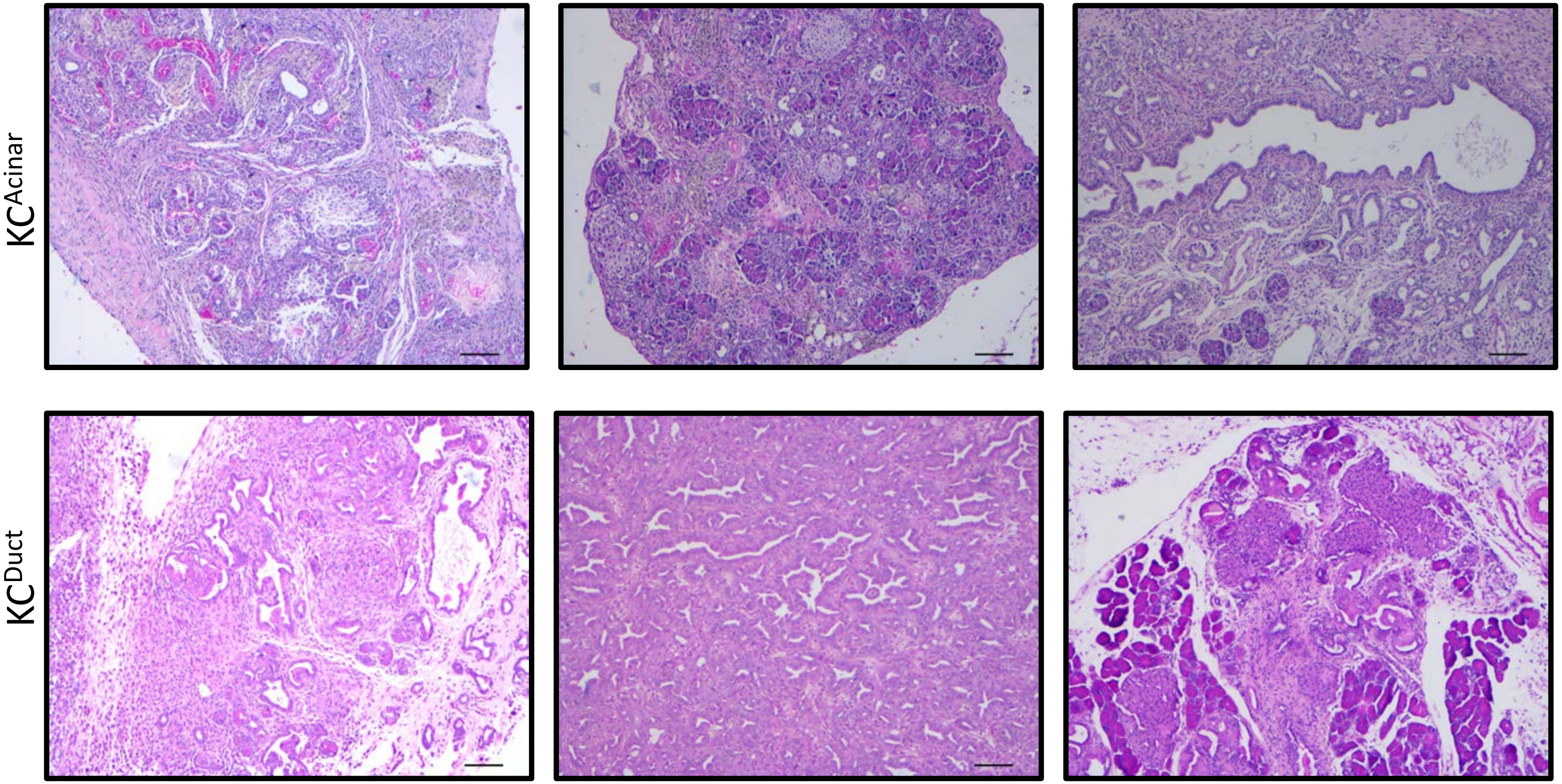


**Supplemental Figure 3.** Hematoxylin and Eosin staining of regions used for IMC experiment.

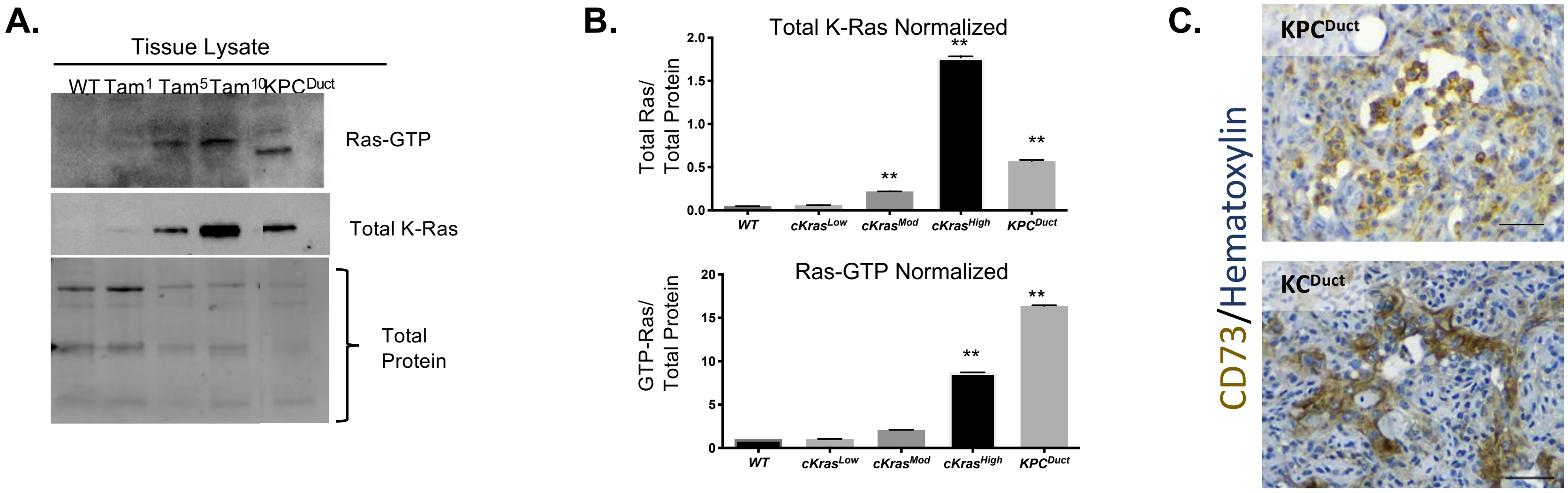


**Supplemental Figure 4. Oncogenic KRAS expression in pancreatic ducts increases levels of CD73. A)** Lysates from wild-type pancreas (Lane1), *Tam1* , *Tam5* and *Tam10* murine pancreas (lanes 2,3,4) were affinity precipitated with Raf RBD agarose and subjected to immunoblot analysis with anti-Ras antibody. Quantification of Ras-GTP binding to total protein reveals. **B**) Quantification of Total Ras protein to total protein confirmed a significant increase in Ras expression with increased tamoxifen dosage. **C)**Immunohistochemistry for CD73 in $KPC^{Duct}$ and $KC^{Duct}$ pancreata show epithelial specific staining for CD73.

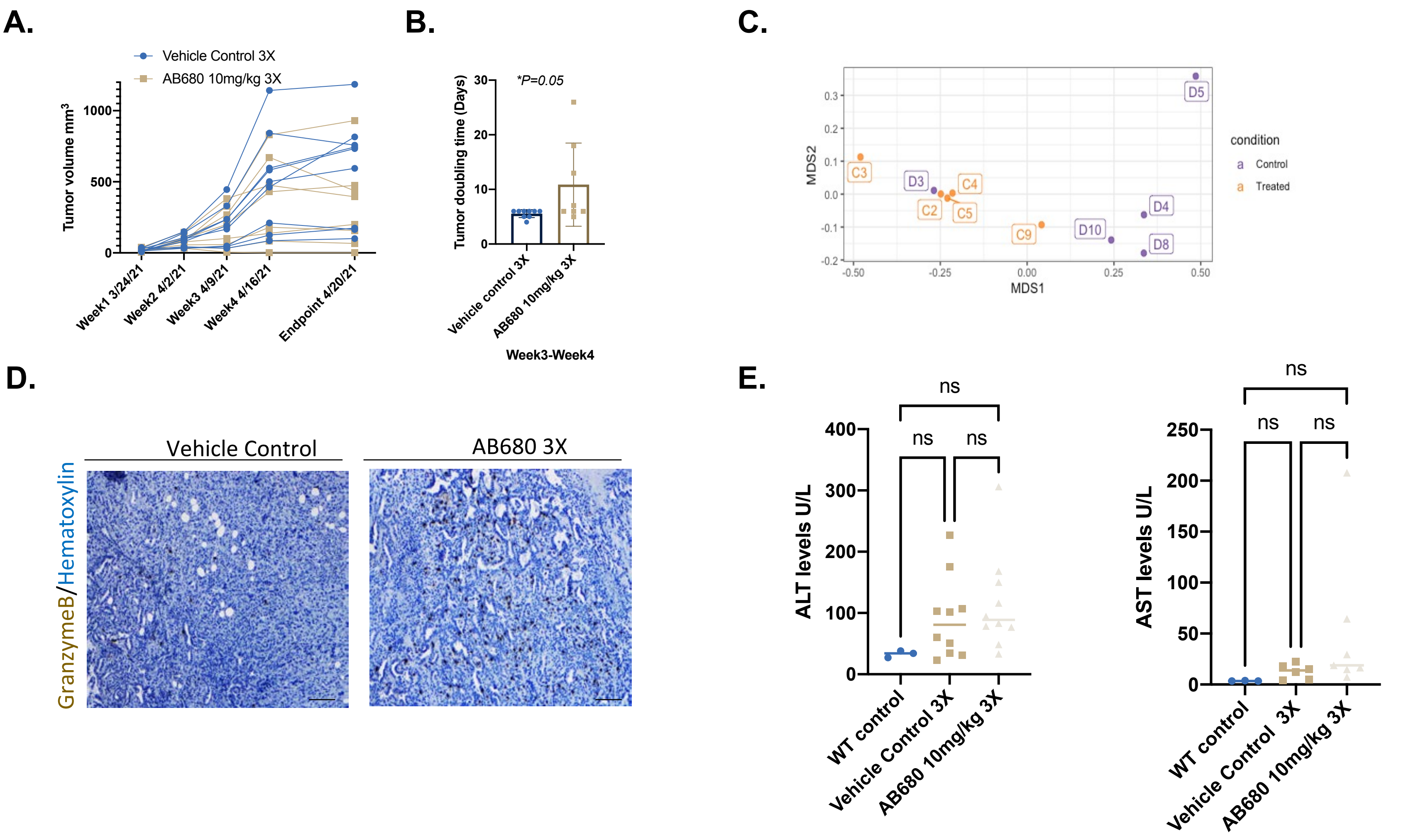


**Supplemental Figure 5. Inhibition of CD73 using AB680 significantly increases anti-tumor immunity and is not associated with hepatic toxicity. A)** AB680 treatment significantly decreased KPC subcutaneous growth rates and **B)** Graphical representation of subcutaneous tumor growth rates for analysis of individual tumor doubling time (**P<0.05*), n=6. **C)** Clustering of control and AB680 treated samples for CyTOF experiment. **D**) Granzyme B IHC analysis in tumors from control and AB680 treated mice. **E**) No significant difference in serum ALT or AST analysis in AB680 treated mice. A student's t test was used to evaluate statistics.
